# Predicting and prioritizing community assembly: learning outcomes via experiments

**DOI:** 10.1101/2022.07.07.499099

**Authors:** Benjamin Blonder, Michael H. Lim, Oscar Godoy

## Abstract

Community assembly provides the foundation for applications in biodiversity conservation, climate change, invasion ecology, restoration ecology, and synthetic ecology. Predicting and prioritizing community assembly outcomes remains challenging. We address this challenge via a mechanism-free *LOVE* (Learning Outcomes Via Experiments) approach suitable for cases where little data or knowledge exist: we carry out actions (randomly-sampled combinations of species additions), measure abundance outcomes, and then train a model to predict arbitrary outcomes of actions, or prioritize actions that would yield the most desirable outcomes. When trained on <100 randomly-selected actions, LOVE predicts outcomes with 2-5% error across datasets, and prioritizes actions for maximizing richness, maximizing abundance, or minimizing abundances of unwanted species, with 94-99% true positive rate and 12-83% true negative rate across tasks. LOVE complements existing approaches for community ecology by providing a foundation for additional mechanism-first study, and may help address numerous ecological applications.

## Introduction

There has been a focus in community ecology on understanding community assembly and coexistence mechanisms (Chesson 2000; Letten *et al*. 2017; Ellner *et al*. 2019). However, predicting and prioritizing community assembly outcomes (Allen-Perkins *et al*. 2023; Houlahan *et al*. 2017; Keddy 1992; Laughlin & Laughlin 2013) is also relevant to applied challenges. Applications include restoration (Palmer *et al*. 1997; Wainwright *et al*. 2018), control or screening of invasive species (Gallien & Carboni 2017; Shea & Chesson 2002), disease ecology (Johnson *et al*. 2015), agriculture (Malézieux 2012; Vandermeer 1995), microbiome engineering and synthetic ecology (Clark *et al*. 2021; Lindemann *et al*. 2016; Nalley *et al*. 2014), and gut microbiome health (Widder *et al*. 2016). Here we focus on advancing these applications when mechanistic insight or data are limited.

We define an outcome as the abundance of species present in a community after a certain amount of time (**Figure 1a**). The outcome does not have to represent stable coexistence (Chesson 2000), but could. In *prediction*, we assume a community is in an initial state *S*_*initial*_ (defined as the abundances of each species present or absent), and that an action (‘experiment’) *A* occurs (e.g., adding a species); then we predict the final state *S*_*outcome*_ (outcome) (**Figure 1b**). In *prioritization*, we also assume *S*_*initial*_ and indicate a desired *S*_*outcome*_; we then determine which *A* should be implemented to yield *S*_*final*_ (**Figure 1c**). That is, we find the action is most likely to yield a desired outcome. Under this framework, effective prediction would enable effective prioritization. If the desirability of an outcome can be estimated, then prioritization proceeds by first, predicting outcomes over all experimental actions; second, enumerating the desirability for each; then third, identifying which action(s) would yield the highest desirability.

**Figure 1.**
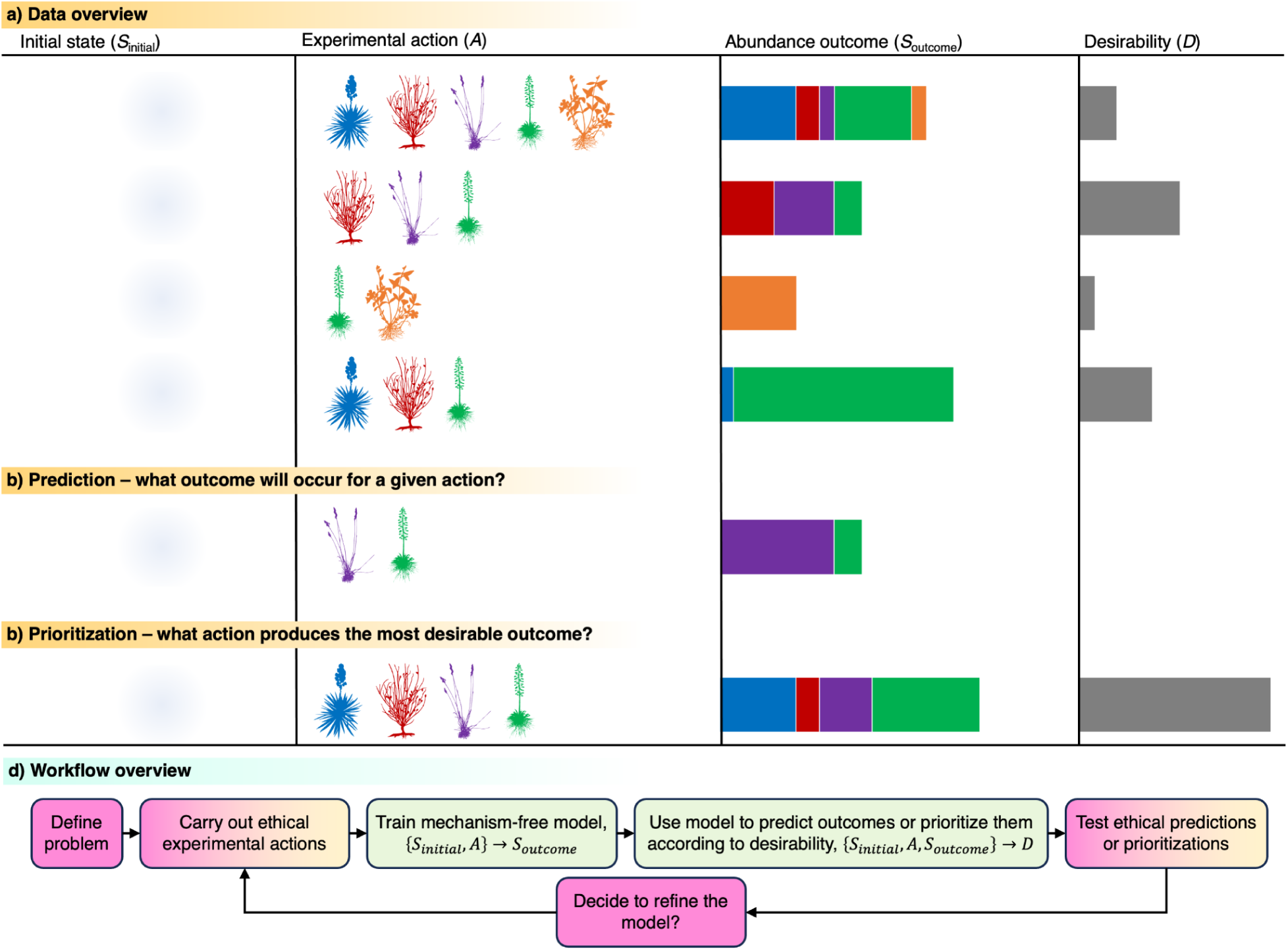
**(a)** Overview of the datasets used by *LOVE*. A community is first observed in an initial state (S_initial_), here assumed to be empty (shadowed region). An experimental action (A) is then taken, here representing a species addition (colored species silhouettes). After some time has passed, an abundance outcome is observed (S_final_), here with bars representing abundances with the same colors as silhouettes. The desirability (D) of the outcome can also be independently determined by humans. **(b)** In prediction, the mechanism-free model is used to determine the outcome of proposed actions. **(c)** In prioritization, the mechanism-free model is used to determine best action(s) within the potential action space that maximize(s) desirability. **(d)** Overview of the inference procedure for *LOVE*. Magenta steps indicate those that require human decision-making; yellow steps indicate those that require experimental work with real organisms; green steps those that require modeling only.

Some approaches to prediction rely on temporal dynamics. Fitting parametric models to time series data (e.g., the generalized Lotka-Volterra (‘GLV’) model (Bucci *et al*. 2016; Stein *et al*. 2013; Ushio *et al*. 2018) is limited by the need to identify the mechanistic processes to include in the model, and by the need for long datasets, which are hard to obtain especially for long-lived organisms. Alternative approaches to parameterize these models through assembling low-richness communities (e.g. singlets and pairs of species across a density gradient), e.g. (Kraft *et al*. 2015; Levine & HilleRisLambers 2009; Vandermeer 1969), or high-richness ‘dropout’ communities (Bai *et al*. 2022; Carlström *et al*. 2019) neglect higher-order species interactions (Mayfield & Stouffer 2017; Pistón *et al*. 2019). Fitting non-parametric forecasting models (Perretti *et al*. 2013; Ye *et al*. 2015) is also possible and avoids mechanism uncertainty. However, these methods require longer time series than typically available (Chang *et al*. 2017) as do other machine learning methods (Baranwal *et al*. 2021; Clark *et al*. 2021; Kong *et al*. 2020; Rammer & Seidl 2019), e.g. >37,000 observations for (Civantos-Gómez *et al*. 2021).

The limitations of these approaches to prediction and prioritization may be overcome if outcomes, rather than temporal dynamics, are of interest. This outcome-focused approach would reduce understanding of community dynamics but potentially have more tractable data requirements, and help when mechanistic insight is not yet available. Several mechanism-free approaches have been developed. For example, studies of observational species co-occurrence outcomes have yielded checkerboard ‘assembly rules’ (Diamond 1975) and joint species distribution models (Pollock *et al*. 2014). However, these methods make strong linearity assumptions or conflate environmental factors with species interactions (Blanchet *et al*. 2020; Connor *et al*. 2013). Studies of experimental co-occurrence outcomes have yielded matrix pseudo-inversion (Maynard *et al*. 2020) or compressive sensing (Arya *et al*. 2023) methods, which are successful primarily when higher-order species interactions are rare. Machine learning has been applied to the design of synthetic microbial communities with more flexibility (Baranwal *et al*. 2022; Chang *et al*. 2021; Clark *et al*. 2021; Connors *et al*. 2023; Lindemann *et al*. 2016; Pacheco & Segrè 2021). Restoration and agriculture applications exist (Fremout *et al*. 2022; Hou *et al*. 2022; Laughlin 2014), but with simpler algorithms and limited consideration of species interactions.

Several conceptual questions around mechanism-free prediction and prioritization exist: (**1**) How does prediction skill for a mechanism-free approach compare to a mechanistic approach? Augmenting a mechanism-free approach with information from expert knowledge or partially-correct mechanisms (e.g., a GLV model) might enhance a mechanism-free model. **(2)** How much training data are needed to reach an acceptable skill level? (**3**) Does the experimental design for gathering training data matter? Many studies have focused on pairwise assembly experiments, but other experimental designs exist, e.g. randomly selected experiments, or active learning (sequential design of experiments). **(4)** What properties of a dataset make it suitable for mechanism-free prediction, e.g., the strength and sparsity of species interactions? (**5**) Which types of prioritization tasks are tractable? **(6)** What properties of a dataset make it suitable for prioritization?

We address these questions using a prediction and prioritization approach called *LOVE* (Learning Outcomes Via Experiments) applied to seven community assembly datasets. We also discuss the practical and ethical considerations relevant to applied ecology challenges.

## Methods

### Concepts

The overall *LOVE* workflow is (**Figure 1d**): (1) define a problem with relevant people, (2) carry out a set of ethical experimental actions and then wait, (3) use the outcomes to train a mechanism-free model; (4) use the model to predict outcomes and/or prioritize actions that would yield the desired outcome; (5) after ethics assessment, test predictions or prioritizations; (6) potentially refine the model with more data. Mathematical components are below; ethical components, in the Discussion.

LOVE approximates a function *f*:{*S*_*initial*_, *A*} → *S*_*outcome*_ (**Figure 1b**). This is a surrogate modeling problem (Forrester *et al*. 2008; Gramacy 2020). In the community assembly problem considered here, we assume that *n* is the species richness of the regional pool, that *S*_*initial*_ ∈ ℜ^*n*^ which describes the abundance of species, is empty; that *A* ∈ {0,1}^*n*^ describes an action (species addition) carried out a common environment, with 0 indicating absence and 1 presence for each species; and that *S*_*initial*_ ∈ ℜ^*n*^ is the abundances of species in the outcome. There are *2*^*n*^ possible unique actions, but an actual study might replicate the same experimental action multiple times with varying outcomes (e.g., due to stochasticity, or uneven success implementing the action) (**Table 1**, **Text S1**).

**Table 1.** Summary of datasets used in this study. The number of experiments indicates the total number of experimental actions available in the dataset; unique experiment numbers may be lower if experiments have been replicated. The number of excluded experiments indicates cases omitted from training due to outlier abundance values. The number of possible experiments is equal to the cardinality of the action space. More detail on dataset provenance and preprocessing is provided in **Text S2**.

| Dataset | *Taxa* | *Provenance* | *Citation* | # of experiments | # of unique experiments | Outcome mean abundance (95% quantile) | # of possible experiments |
| --- | --- | --- | --- | --- | --- | --- | --- |
| Annual plant | California annual plants | Simulations from nonlinear competition / seedbank model | (Godoy *et al.* 2014, 2017) | 262144 | 262144 | 4321.1 | 262144 |
| Cedar Creek | North American prairie plants | Experimental sowing in natural environments | (Tilman *et al.* 2001, 2012) | 154 | 132 | 50.2 | 262144 |
| Fly gut | Bacteria in fly gut | Experimental inoculations of germ-free flies | (Gould *et al.* 2018) | 1536 | 32 | 537000 | 32 |
| Human gut | Bacteria in human gut | Simulations from GLV competition model | (Venturelli *et al.* 2018) | 4096 | 4096 | 0.6 | 4096 |
| Mouse gut | Bacteria in mouse gut | Simulations from GLV competition model | (Buffie *et al.* 2012; Stein *et al.* 2013) | 2048 | 2048 | 13.3 | 2048 |
| Soil bacteria | Bacteria in soil | Experimental assembly in microcosms | (Friedman *et al.* 2017) | 570 | 101 | 0.1 | 256 |
| SORTIE-ND | Eastern North American hardwood<br>trees | Simulations of forest | (Pacala *et al.* 1996) | 1536 | 512 | 195 | 512 |

### Mechanism-free methods for learning *f*

#### Naïve

In a null approach, we obtained outcome predictions using a heuristic. We assumed the abundance of each species in *S*_*outcome*_ was equal to its mean abundance in the training data, elementwise-multiplied by *S*_*initial*_.

#### Random forest

Random forest classifiers were used because they allow for nonlinear and multiple interactions among predictors, often avoid overfitting, and are suitable for sparse datasets (Breiman 2001). Models were trained using the *randomForestSRC* R package (Ishwaran & Kogalur 2019) (version 2.12.1). Models were fit using *num.trees*=500, *mtry*=*ceiling(sqrt(n))*, and *nodesize*=5. To reduce the impact of zero-inflation and skewness, abundances were binned into ten classes, comprising 0, eight quantiles of the non-zero abundance values (over the whole dataset), and the maximum abundance. Predicted class values were transformed back into abundances as either 0 or the bin-mean value. The number of bins did not have a large impact on results (not shown).

#### Sequential random forest

We assessed the value of active learning, where training cases are selected sequentially to maximize information gain. We developed a sequential random forest method adapted from (Gu *et al*. 2015) that selects action vectors that would yield the greatest information gain. We performed 10 active learning iterations, sequentially collecting an additional 1/10th of the data in each iteration to create a full training dataset. For each iteration of active learning, we selected the actions with the highest score, with score defined as the sum of:

- Uncertainty: the variance of the bootstrap predictions for the candidate action, for 5 bootstrap samples of the data collected until that step.
- Diversity: the sum of the Hamming (L^1^) distance between the candidate action and the 10 closest action vectors within the training set.
- Density: the Hamming distance between the candidate action vector and other unsampled action vectors.

#### GLV model

We also compared our method to EPICS, a GLV fit to outcome data (Ansari *et al*. 2021). To enable EPICS to handle missing data and duplicate training data points common to our datasets, we developed a modified version, gEPICS. In the original approach, their 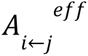 matrix was calculated by solving 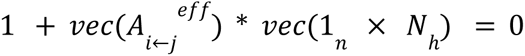 where *N_h_* is their notation for species abundance. By calculating the matrix inverse 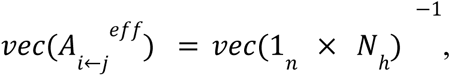 which is guaranteed to exist in the original problem formulation, they obtained the outcome abundance. In gEPICS, we instead calculated the generalized Penrose pseudoinverse (†),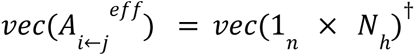. With the estimated 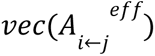) matrix, we then performed estimation of the experimental outcome by calculating the generalized analog of the GLV nullcline solution, 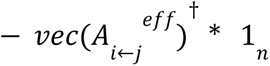. We replaced any negative predicted values with 0.

#### Random forest + GLV residuals

We developed a residual learning approach (building on successes in image recognition (He *et al*. 2016)), combining model components (gEPICS) and residual effects that cannot be explained by GLV (random forest). First, we fit the gEPICS model on the dataset. Second, we predicted the abundances with the fitted GLV model and obtained residuals. Third, we fit a random forest model on the residuals with no abundance binning. For final outcome predictions, we summed the gEPICS and random forest prediction values.

#### Random forest + GLV features

We gave the random forest model additional information from a GLV model. We used the random forest method, but with the input variables including the experimental actions and also the GLV prediction values obtained by fitting a gEPICS model.

### Experimental designs

#### Low richness

There are 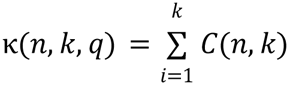 possible assemblages with richness ≤*k*, where C(*n*,*k*) indicates the binomial coefficient. We selected a random set of cases for training, selecting only among assemblages with *k*=2, or *k*=3 pairs and triplets, named as the *low-2* and *low-3* experimental designs. No additional cases are selected after all pairs and triplets are exhausted.

#### High richness

There are also κ(*n*, *k*) possible assemblages with richness ≥*k*. We selected a random set of cases for training, selecting only among assemblages with *k*=*n*-1, *k*=*n*-2 (single or double dropouts, named as the *high-1* and *high-2* experimental designs). No additional cases were selected after all single and double dropouts are exhausted.

#### Mixed richness

we selected a random set of cases from each dataset for training independent of richness (named as the *mixed* experimental design). Because κ(*n*, *k*) is largest at *k*=⌊*n*/2⌋, intermediate richness assemblages are frequently sampled.

#### Prior

we selected states that are either singlets (only one species present), or leave one out (all but one species present). Further cases are sampled according to the *mixed richness* design, mirroring (Ansari *et al*. 2021).

#### Sequential

we sampled initial training data according to the *mixed richness* design, and then add data points in batches according to, and only for, the sequential random forest method.

### Datasets

Seven empirical and empirically parameterized datasets of combinatorial community assembly experiments were used, spanning a range of taxa (**Table 1**, **Text S1**, **Figure S1-S7)**. For datasets generated from a parameterized dynamical model, only the predicted outcomes are used. All datasets were pre-processed to first remove outcome abundances exceeding 10^7^, which arose in a few assemblages within the ‘mouse gut’ dataset, and then were clipped to the (0.005,0.095) quantiles (across all assemblages within each dataset) to avoid outlier overfitting.

### Analyses

Analyses were carried out in a training-testing cycle for each algorithm, experimental design, and sample size. Each analysis was replicated 10 times to capture training case sampling variation. Training-set sample sizes spanned from 10 to 10,000, covering 20 values evenly spaced on logarithmic scale. Analyses were skipped where sample sizes exceeded either the dataset size or the maximum number of samples available for the experimental design. We then compared predicted outcomes to actual outcomes in the test-set assemblages. Scaled error was defined as the mean absolute error (MAE) between the observed and predicted *S*_*outcome*_ scaled by the 95% quantile dataset abundance, treating each experimental action as a replicate.

For Questions 1-3, we plotted marginal predictions for the test-set scaled error rate (*scaled_error*) as a function of the method (*method*), the training sample size (*num_train*), and the experimental design (*experimental_design*), and the dataset. To reduce the high dimensionality of the dataset and reflect a realistic use case, for *method* we used *random forest*; for *num_train*, 89; for experimental design, *mixed*. Because the data have a statistically balanced design, no *post-hoc* model was used.

For Question 4, we plotted the test-set scaled error rate as a function of several dataset properties: whether the dataset was generated from real experiments or from dynamical model simulations (*type*), the number of species in the regional pool for each dataset (*regional_pool_richness*), the mean number of species gained or lost from the experiment to the outcome (*num_losses_mean)*, and the mean of the skewness of the abundances of species present in the outcome (*abundance_skewness_mean*). We conditioned on values for *method* of *random forest*; for experimental design, *mixed*. Because predictors are potentially correlated, we fit a linear mixed model:

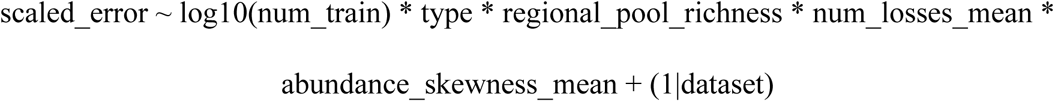

We visualized model predictions using conditional effect plots and summarized fit using Nakagawa’s pseudo-R^2^.

For Question 5, analyses were restricted to the four datasets where the complete set of experimental outcomes were available for validation (*annual plant*, *human gut*, *mouse gut*, *SORTIE-ND*). We prioritized experiments as described below, then compared the prioritized experiments to the actual best experiments using true positive and negative rate metrics. We assumed that there existed a desirability function via a function *g*: _{_*S*_*initial*_, *A*, *S*_*outcome*}_ → *D*. For simplicity, we assume this function is determined entirely by these predictors, in contrast to a more complex approach where *D* is a learned function (Clark *et al*. 2021; Connors *et al*. 2023).

In a ‘remove unwanted’ desirable outcome, we searched for communities that would be invasion-resistant. Desirable outcomes were identified as those where a focal species *i* was present in the experiment and occurred at its 0% quantile abundance in the outcome (0 if ever absent in at least one outcome, or minimum abundance if never absent in any outcome), i.e. *D*_*i*_ = (*S*_*initial*,*i*_ > 0)×(*S*_*outcome*,*i*_ = 0). We repeated this analysis for every species in every dataset.

In a ‘maximize diversity’ desirable outcome, we searched for communities with high biodiversity (Shannon’s index; (Pielou 1966)). We predicted abundance for all non-training set experiments and calculated a predicted H value as the desirability function, i.e. 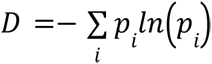 where 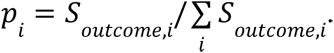. Desirable cases were flagged as those with a D above the 95% quantile D value actually observed in all assemblages (in a real-world use, this quantile threshold’s value would be unknown *a priori*, but a known threshold value for D could be specified).

In a ‘maximize abundance’ desirable outcome, we searched for communities with high summed abundance across all species, i.e. 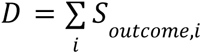. Desirable cases were flagged as those with a D above the 95% quantile D value actually observed in all assemblages.

Data for prioritization come from a *random forest* method, a *mixed richness* experimental design, and a num_train of either 89 or 264. Analyses were replicated across 10 sampled training datasets. We then summarized the true positive and true negative rates of the prioritized experiments relative to the actual best experiments. We also visualized the similarity between the prioritized experiments and outcomes relative to their actual values, using heatmaps with cases hierarchically clustered by Euclidean distance. We additionally carried out a principal component analysis of the outcome abundance space, then visualized the distribution of classifications for each experiment within this space.

For Question 6, we plotted the true negative rate of the prioritization for each task as a function of *regional_pool_richness*, *num_losses_mean*, and *abundance_skewness_mean*. We conditioned on values for *method* of *random forest*; for experimental design, *mixed* and fit a linear mixed model:

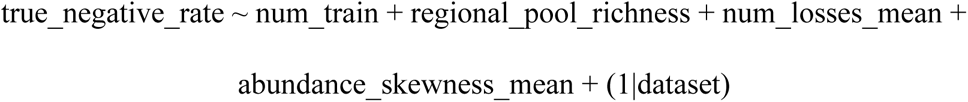

Fixed effect interactions were not included due to the sample size. In the removal model, a random intercept for removed species was also included. We visualized model predictions using conditional effect plots and summarized fit using Nakagawa’s pseudo-R^2^.

### Data availability statement

Processed datasets and code are available at https://github.com/bblonder/coexistence_LOVE.^1^

## Results

### Question 1 - value of mechanism-free prediction and mechanism

The mechanism-free methods performed as well or better than a mechanistic method at predicting abundance in experimental outcomes across all datasets (**Figure 2a**). The naïve method obtains an error rate of 10-50% depending on the dataset. The GLV model often had error rates substantially higher than this baseline, and required large numbers of training experiments (>500 depending on dataset) to reach lower error rates. This is notable as several datasets are from simulations of a GLV model. Providing the random forest method with additional residuals from a GLV fit (i.e. a residual learning approach) had no effect, while the random forest method on those residuals directly was worse that the GLV fit.

**Figure 2.**
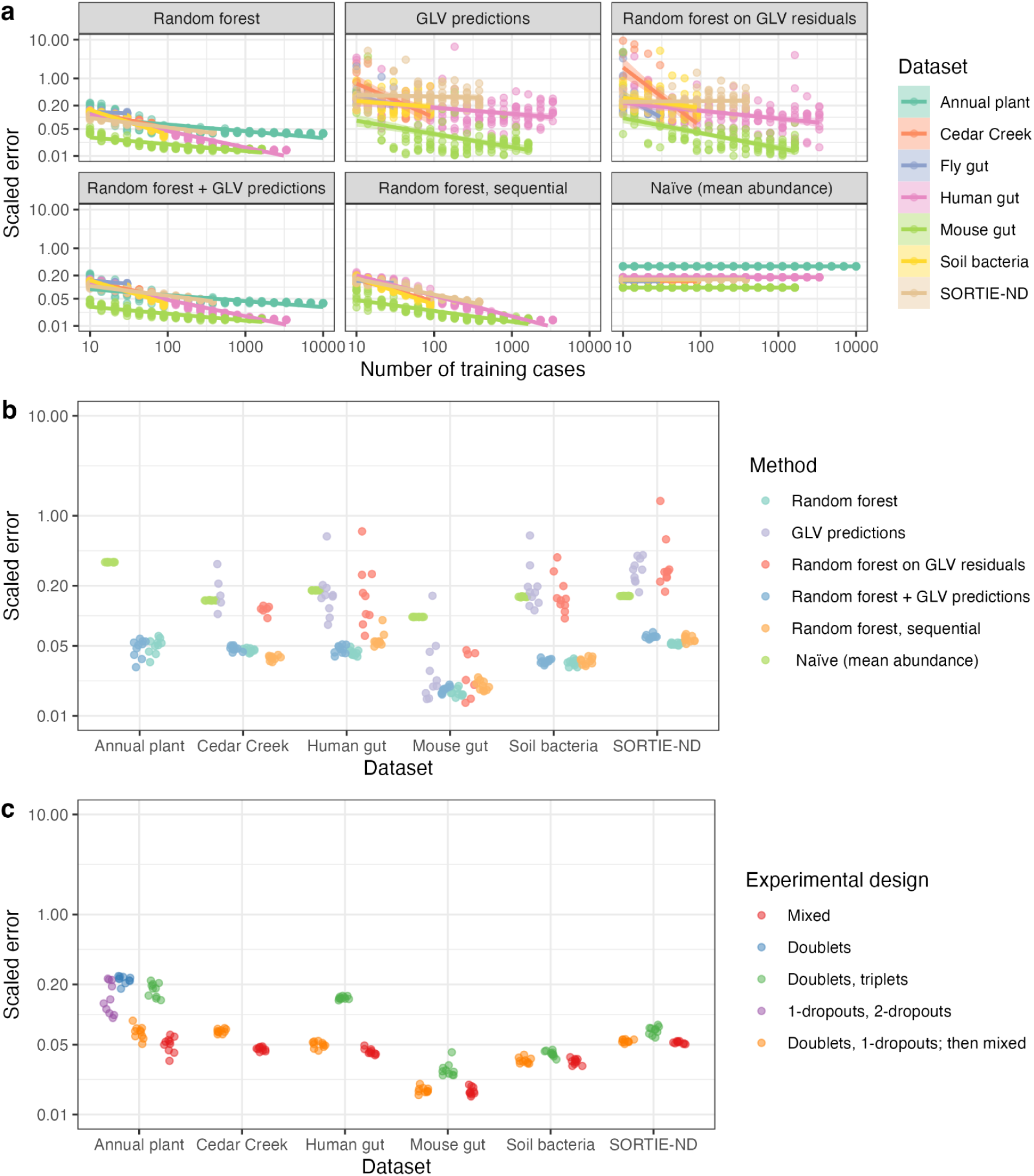
Comparison of abundance prediction skill in several scenarios. All panels’ y-axis indicates the mean absolute error in abundance scaled by the datasets’s 95% quantile abundance; lower values indicate better prediction skill. Results from ten training replicates are shown as points. The y-axis scale is log-transformed. **(a)** Comparison of methods, breaking out the effect of the number of training experiments, conditioning on a mixed richness experimental design. **(b)** Comparison of methods, conditioning on a mixed richness experimental design and 89 training experiments. **(c)** Comparison of experimental designs, conditioning on a random forest method and 89 training experiments. In panels b and c, some designs and datasets are not shown due to an insufficient number of training experiments.

When comparing the methods at a plausible number of training experiments (89), the baseline random forest and the sequential random forest had lowest error rates (**Figure 2b**).

### Question 2 - number of experiments required

The mechanism-free methods yielded error rates below the naïve baseline typically by ∼50 training experiments, and continued to improve in skill with more experiments (**Figure 2a**). The mean scaled error rate dropped to 2-5% across datasets with <100 experiments (**Table S2**). The sequential random forest was only slightly more efficient at learning from training experiments than the random forest.

The structure of abundance error is shown for a random forest method, a mixed richness experimental design, and 89 training experiments (**Figure S8)**. Errors were generally unbiased, though a small number of species were consistently unpredictable, with lower or variable abundances than observed. False prediction of absence was the main systematic error. All of these issues become unimportant at larger sample sizes (e.g, 264 training experiments; **Figure S9**).

### Question 3 - experimental design

The lowest error rates were obtained using a mixed richness experimental design, for all datasets (**Figure 2c**). The design of sampling doublets and 1-dropouts before proceeding to mixed richness sampling had similar but slightly worse performance. The doublet, triplet, and dropout experimental designs had error rates up to four times higher than mixed richness sampling.

### Question 4 - dataset properties predicting prediction skill

Some datasets had consistently higher error rates. Some of this variation was explainable by a *post-hoc* mixed model, conditioned on a *random forest* method, a *mixed* experimental design, and 89 training experiments (**Figure 3**). This model had a marginal R^2^ of 64% and a conditional R^2^ of 90%. Higher error occurred for datasets with higher species richness, lower number of species lost, greater outcome abundance mean skewness, and for empirical origins.

**Figure 3.**
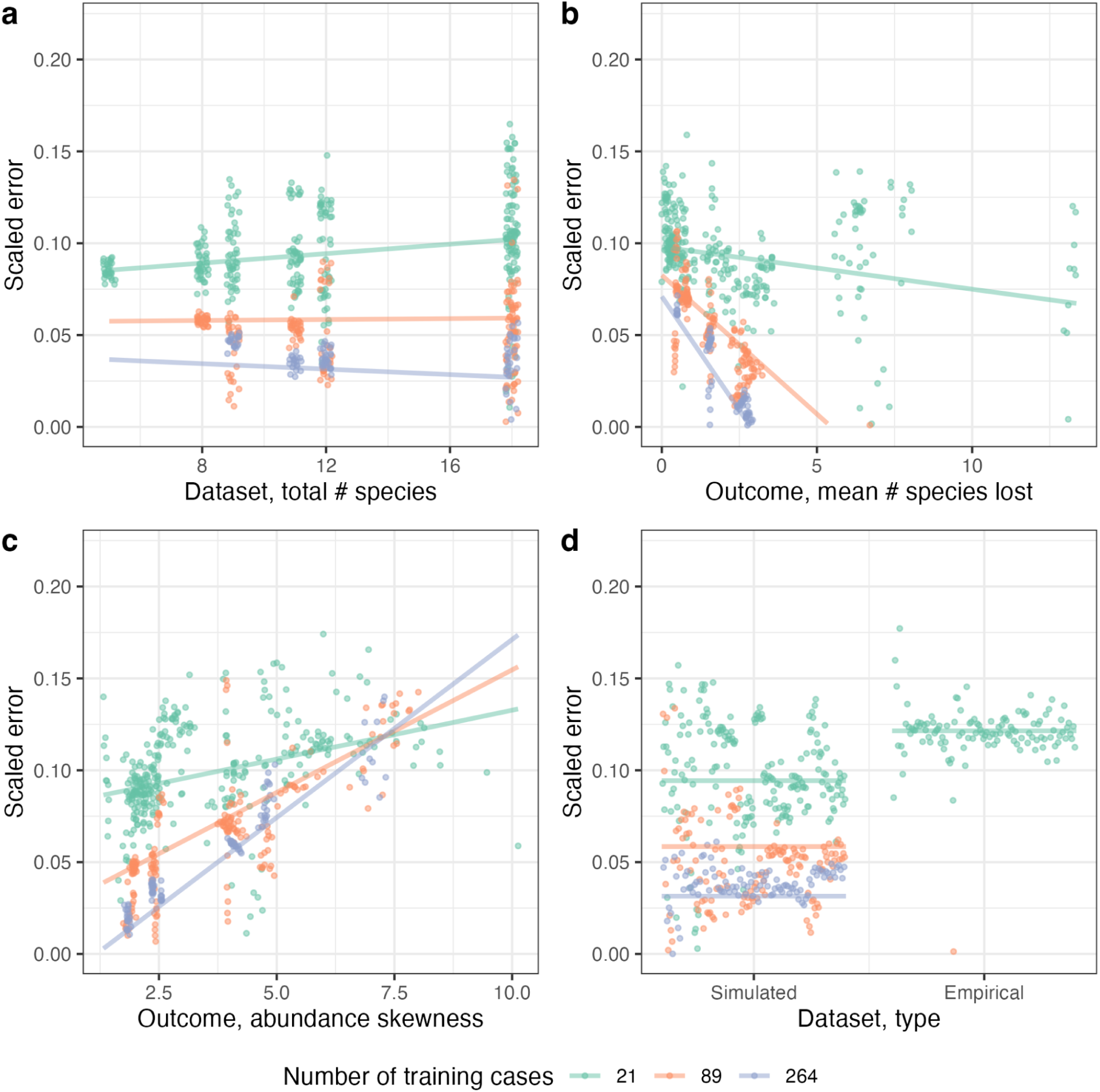
The effect of dataset properties on scaled error using a post-hoc linear mixed model. Predictions are for a random forest method and a mixed richness experimental design. Dots indicate results for each training replicate; lines indicate predicted conditional effects. Panels show the effect of **(a)** species richness of the dataset, **(b)** mean number of species lost (i.e. present in experiment, absent in outcome) in training data, **(c)** mean skewness of outcome abundances in training data, and **(d)** whether the outcomes are from empirical experiments or simulated experiments from a hidden dynamical model.

### Question 5 - tractable prioritization tasks

Skill varied with each prioritization task. When considering a random forest method, a mixed richness experimental design, and 89 training experiments, mean true positive rate varied from 94-99% and true negative rate from 12-84% across tasks (**Table S3**).

For the removal of unwanted species (**Figure 4a**), true positive and true negative rates were >75% in most datasets for most species. However, in each dataset, there were a small number of species for which the true negative rate was always <20%; this likely reflects an absence of training data covering certain species combinations.

**Figure 4.**
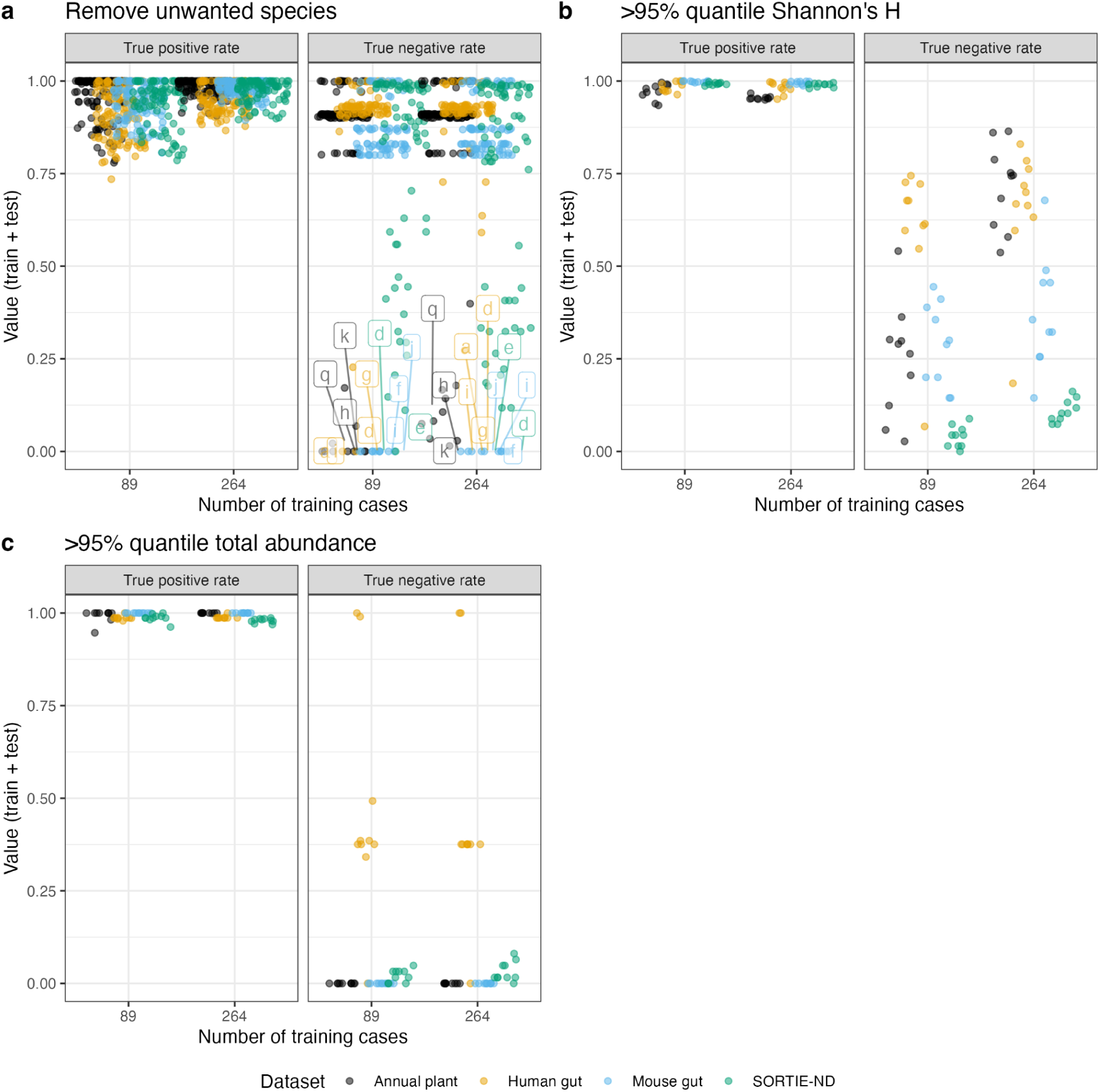
Skill at prioritizing experiments for three prioritization tasks: **(a)** removing an unwanted species, **(b)** obtaining high Shannon’s H, and **(c)** obtaining high abundance. Dots indicate results for each training sample and are colored by dataset. Prioritizations are for a random forest method, a mixed richness experimental design, and 89 training experiments. In panel a, more dots are present because the analysis is repeated for each potential species to remove; letters are shown only for species in which prioritizations for all replicates had <20% true negative rate. Species names for each alphabetical species code are in **Table S1**.

For obtaining high diversity (**Figure 5b**), true positive rate was >80% in all datasets, while true negative rate varied from 0-75%, with some datasets (e.g., human gut) consistently performing well and other datasets (e.g., SORTIE-ND) consistently performing poorly. Somewhat worse results were found for the obtaining of high abundance (**Figure 4c**).

**Figure 5.**
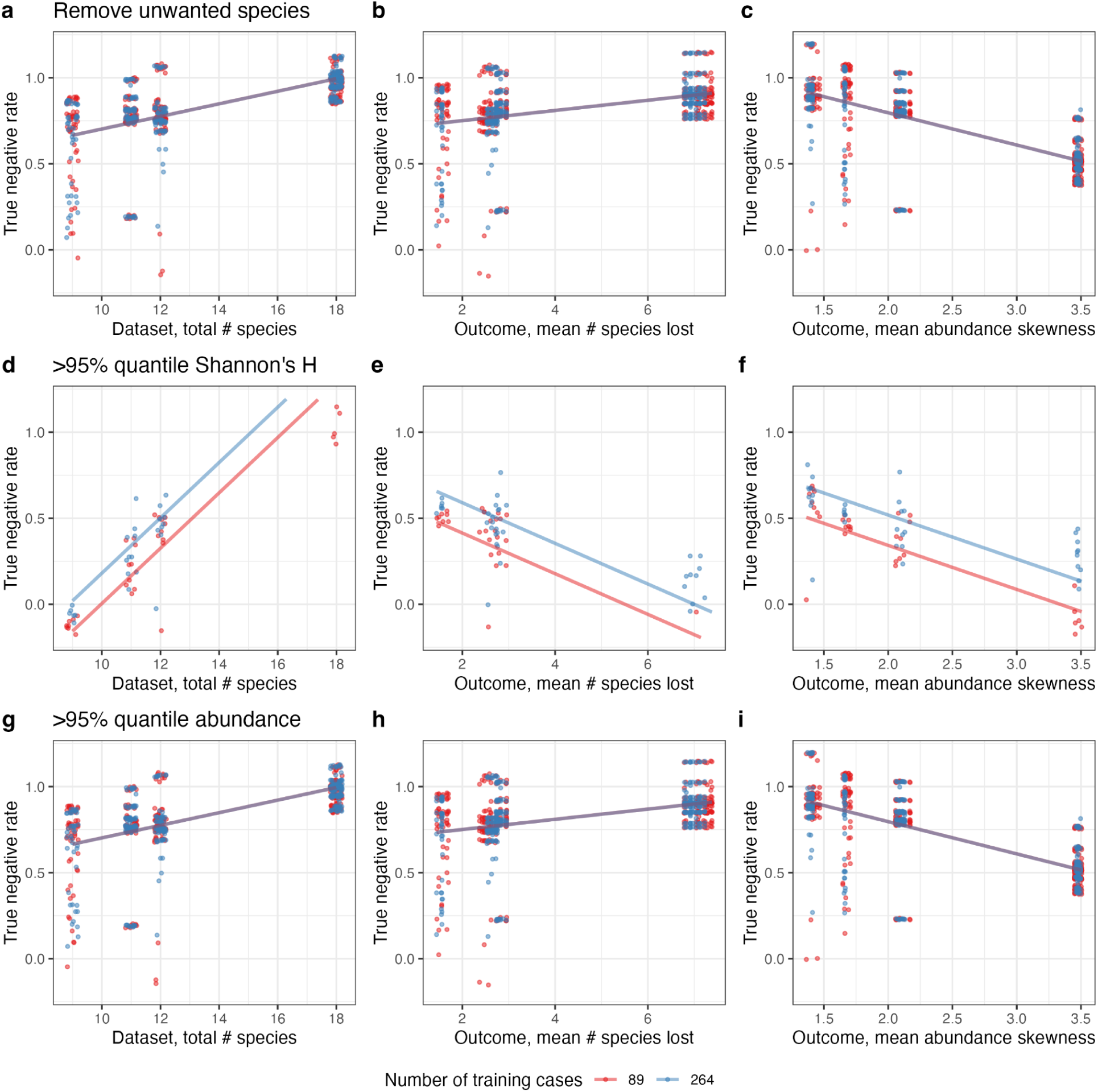
The effect of dataset properties on prioritization true negative rate using a post-hoc linear mixed model. Rows show each of the three prioritization tasks. Predictions are for a random forest method and a mixed richness experimental design. Dots indicate results for each training replicate; lines indicate predicted conditional effects. Panels show the effect of **(a,d,g)** species richness of the dataset, **(b,e,h)** mean number of species lost (i.e. present in experiment, absent in outcome) in training data, and **(c,f,i)** mean skewness of outcome abundances in training data.

When increasing the training sample size to 264, improvement in true negative rate sometimes occurred. For the maximizing Shannon’s H task, 10-50% improvement was possible depending on the dataset. However limited improvement was obtained for the removal and maximizing total abundance tasks.

The structure of prioritization error is shown for the removal (**Figure S10**), diversity (**Figure S11**), and abundance (**Figure S12**) tasks. In the removal and maximizing Shannon’s H tasks, the distribution of prioritized experiments and the actual best experiments is similar. The prioritized experiments typically leverage species that are correctly predicted at high abundances. Errors occur when experiments fail to include species incorrectly predicted to occur at low abundances. In the maximizing total abundance task, the distribution of prioritized experiments and the actual best experiments shows low similarity, consistent with low true negative rate.

The distribution of error types in abundance outcome space depended on the task and dataset (removal, **Figure S13**; maximizing Shannon’s H, **Figure S14**; maximizing total abundance, **Figure S15**). False negatives and false positive errors consistently occurred in different parts of the abundance outcome space.

### Question 6 - dataset properties predicting prioritization skill

Several predictors explained variation in prioritization skill (**Figure 5**). For the removal task, the true negative rate increased with higher species richness, higher mean number of species lost, and lower mean abundance outcome skewness (p<10^-4^). However this model had a marginal R^2^ of 0.07 and conditional R^2^ of 0.63. For the maximizing Shannon’s H task, true negative rate increased with higher species richness (p<0.05), lower mean number of species lost, and lower mean abundance outcome skewness. This model had a marginal R^2^ of 0.49 and conditional R^2^ of 0.66. For the maximizing total abundance task, true negative rate increased with higher species richness, higher mean number of species lost, and lower mean abundance outcome skewness (p<10^-3^). This model had a marginal R^2^ of 0.62 and conditional R^2^ of 0.62.

## Discussion

Outcome prediction can be successful without understanding dynamics or community assembly processes. Mechanism-free approaches are complementary to other mechanism-first or generality-oriented approaches (Evans *et al*. 2013; Levins 1966). They avoid the complexity of community dynamics and the limitations of mechanistic assumptions, e.g. competition (Simha *et al*. 2022). They provide a useful first step, with low data requirements, towards further mechanistic understanding.

Simple algorithms and sparse datasets (mixed richness sampling of training data, a random forest algorithm, and less than 100 experimental actions for training) yielded acceptable results. For prediction, <5% abundance error was obtained. For prioritization, high true positive rate and variable true negative rate was obtained. True negative rates were typically above 20%, indicating that at least 1 in 5 prioritized experiments would lead to the desired outcome, far better than what random selection of experiments would yield.

Mixed sampling of experimental actions provided more information per experiment than other designs, due to the multiple species combinations that are simultaneously explored. Additionally, such experiments can be carried out in parallel, limiting the total time needed for the approach. In contrast, dropout communities alone are less useful - multiple levels of dropouts are required to resolve complex species interactions, e.g. (Finkel *et al*. 2019). Pair and triplet designs were most successful only when the underlying dynamical model involves purely pairwise interactions. Fractional factorial designs may have higher efficiencies (Gunst & Mason 2009; Santner *et al*. 2003), but may not be optimal if the strength of higher-order species interactions is unknown.

There was substantial variation in skill across the datasets explored. For prediction, smaller state spaces, stronger species interactions, fewer rare species with high abundance, and lower stochasticity all reduce error. For prioritization, large state spaces and few dominant species both reduce error (because even low true positive rates are useful when state spaces are large). However, future studies could identify additional mechanisms that make outcomes more or less predictable. For example, a fully neutral community assembly process would yield random outcomes and high error. It remains unknown how the topology of interaction networks or the nonlinearity of interactions might affect skill.

Guidance from a mechanistic model was not helpful for prediction. Model residuals or model predictions did not improve skill relative to the random forest, nor did more complex experimental designs. Simple algorithms for function approximation may already leverage all the information present in the data, consistent with findings from (Arya *et al*. 2023).

### Conceptual considerations

*LOVE* is best used when data are sparse and regional pool richness is high. When *n* is large, *LOVE* requires a small number of experiments relative to the size of the action space and provides a useful approximation. In contrast, when *n* is small, the action space can simply be enumerated via trying all experiments. For example, in the ‘fly gut’ dataset (*n*=5), *LOVE* had low utility because the number of training experiments was close to the number of actual possible experiments. Many currently available datasets do not achieve high coverage of the action space. For example, in the Cedar Creek (USA) and Jena (Europe) biodiversity-ecosystem functioning studies, less than 1% of all possible plant communities were experimentally assembled (Tilman *et al*. 2012; Weisser *et al*. 2017). Shifting to a mixed experimental design for similar future studies would be valuable for applying *LOVE*.

Because of the function approximation approach, there is no ability to extrapolate to novel environmental conditions, actions, or species. However, it should be possible to include environmental conditions as additional dimensions, but training would likely require replicating experiments across an environmental gradient, e.g., (Pennekamp *et al*. 2018). It should also be possible to add trait predictors to augment species identity, which could allow extrapolation of the effects of novel species. However, in species invasion prediction, trait-based approaches have had uneven success (Drenovsky *et al*. 2012; Fournier *et al*. 2019; Thompson & Davis 2011).

The amount of time to wait between the action and the outcome is implicit in *LOVE*. It is assumed to be determined by the investigator’s interests and practical constraints. Sometimes it may be possible to assume an equilibrium has been obtained, and/or that the outcome represents stable coexistence, but not always (e.g. the transient stochastic dynamics in the SORTIE-ND simulation and dataset). That is, *LOVE* makes inferences about persistence, not about stable coexistence; mechanistic approaches are required for the latter.

Methodological improvement may be possible. Zero-inflation can cause challenges. Two-stage models or class weighting approaches can be used to address this in the single-species context but are not feasible in multi-species contexts due to correlations in abundance (especially zeros) across species. We converted abundances to factors, which reduces the impact of zeros, but multivariate hurdle approaches may work better (Kong *et al*. 2020). Additionally, the structure of errors in abundance space for prioritization suggests that an additional model could be coupled to the prediction model to better approximate the *D* function (i.e. learning rather than enumerating prioritization).

More complex community assembly problems could also be studied (Blonder *et al*. 2023). Initial states could be non-empty. Actions could be continuously-valued (*A*∈*R*^*n*^) to reflect variation in the magnitude of a species addition. The dimensionality of *A* could be increased to assess order-of-arrival effects (Fukami 2015; Weidlich *et al*. 2021). The dimensions of S and A could also be increased to allow for environmental covariates, e.g., (Pennekamp *et al*. 2018).

Applications may initially be most successful for short-lived organisms and controlled environments (e.g., microbiomes). Prioritization applications to long-lived organisms may require long waiting times beyond the timeline of decision-making. Similarly, the assumption of a fixed environment may not be valid if temporal environmental change occurs.

A non-sequential approach is most realistic when decision-making timelines are limited and experimental actions take a long time, because sequential methods require multiple iterations. At low numbers of experiments, sequential learning was slightly more data-efficient than non-sequential learning, but required ten times more iterations. While sequential design of experiments (Santner *et al*. 2003) and active learning (Shalev-Shwartz 2012) have many uses, they seem less practical here. Similarly for prioritization, a Bayesian optimization approach requiring multiple iterations to simultaneously learn best actions and outcomes is probably not realistic.

### Ethical considerations

Because many of the candidate applications of *LOVE* are in applied ecology, it is necessary to consider related ethical issues. Many of the potential applications involve making predictions that involve high-risk species (e.g. an invader). While simulations and initial experiments can build statistical confidence that experiments will yield the desired outcome, there is no way to guarantee it. Outcomes may actually occur that are undesirable, and that may not be recoverable. Additionally, the algorithm may prioritize unsafe experiments, i.e. with transient dynamics that pass through dangerous community states (Aswani *et al*. 2013), or yielding the loss of culturally important species. A healthy respect for ecological complexity (Lawton 1999; Simberloff 2004) and unforeseen consequences (Crichton 1991) is prudent, as is follow-up mechanistic study.

*LOVE* also enables the possibility of dual use, i.e. adversarial applications. It could be possible to discover and then implement actions that *intentionally* cause dangerous outcomes, e.g. rapid loss of biodiversity or introductions of invasive species. For example, drug discovery algorithms (Gupta *et al*. 2021) intended for health applications also can identify novel molecules that are more lethal than known nerve agents (Urbina *et al*. 2022). Dangerous communities may be assemblable that do not exist naturally.

Potential *LOVE* applications must also consider whether they take a technocratic and algorithm-first approach to mediating relationships between people and nature. Such framings could be harmful because they would de-legitimize the value of traditional and expert knowledge, and could support the legacy of colonialism and white supremacy in ecology (Chapman *et al*. 2021; Wyborn & Evans 2021). Real applications of LOVE should include engaging relevant communities, and consideration of the unintended consequences of algorithm deployment.

## Author contributions

BB conceived the idea, processed the datasets, wrote initial code, and drafted the manuscript. ML revised and expanded code and implemented algorithms. OG contributed a dataset and contributed substantially to the manuscript.

## Data accessibility statement

All data re-used in this study are publicly available. Pre-processed data and statistical analysis code are available at https://github.com/bblonder/coexistence_love and will be archived upon acceptance.

## Acknowledgments

Daniel Maynard’s work was the inspiration for this study. Pierre Gaüzère, Lars Iversen, and Courtenay Ray guided initial discussions. Carl Boettiger, Erin Carroll, Jashvina Devadoss, Marcus Lapeyrolerie, Ilaíne Matos, Zachary Sunberg, Claire Tomlin, several anonymous reviewers, and others provided feedback on manuscript drafts. Andrew Letten, Daniel Maynard, Brian McGill, William Petry, and William Sharpless provided feedback and provided guidance on datasets. Lora Murphy and Charles Canham provided input on the SORTIE-ND model. Peter Adler, Jonathan Friedman, Alison Gould, Nathan Kraft, Margaret Mayfield, Sara Mitri, Peter Reich, Alejandra Rodriguez Verdugo, Alvaro Sanchez, Serguei Saavedra, Jürg Spaak, and Ophelia Venturelli provided guidance on datasets. The Cedar Creek experiment was supported by NSF via grant DEB-1831944. OG acknowledges financial support provided by the Ministerio de Ciencia, Innovación y Universidades (RYC-2017-23666). ML acknowledges financial support provided by the National Science Foundation (DGE-1752814, DGE-2146752).

## Supporting Information

**Table S1.**
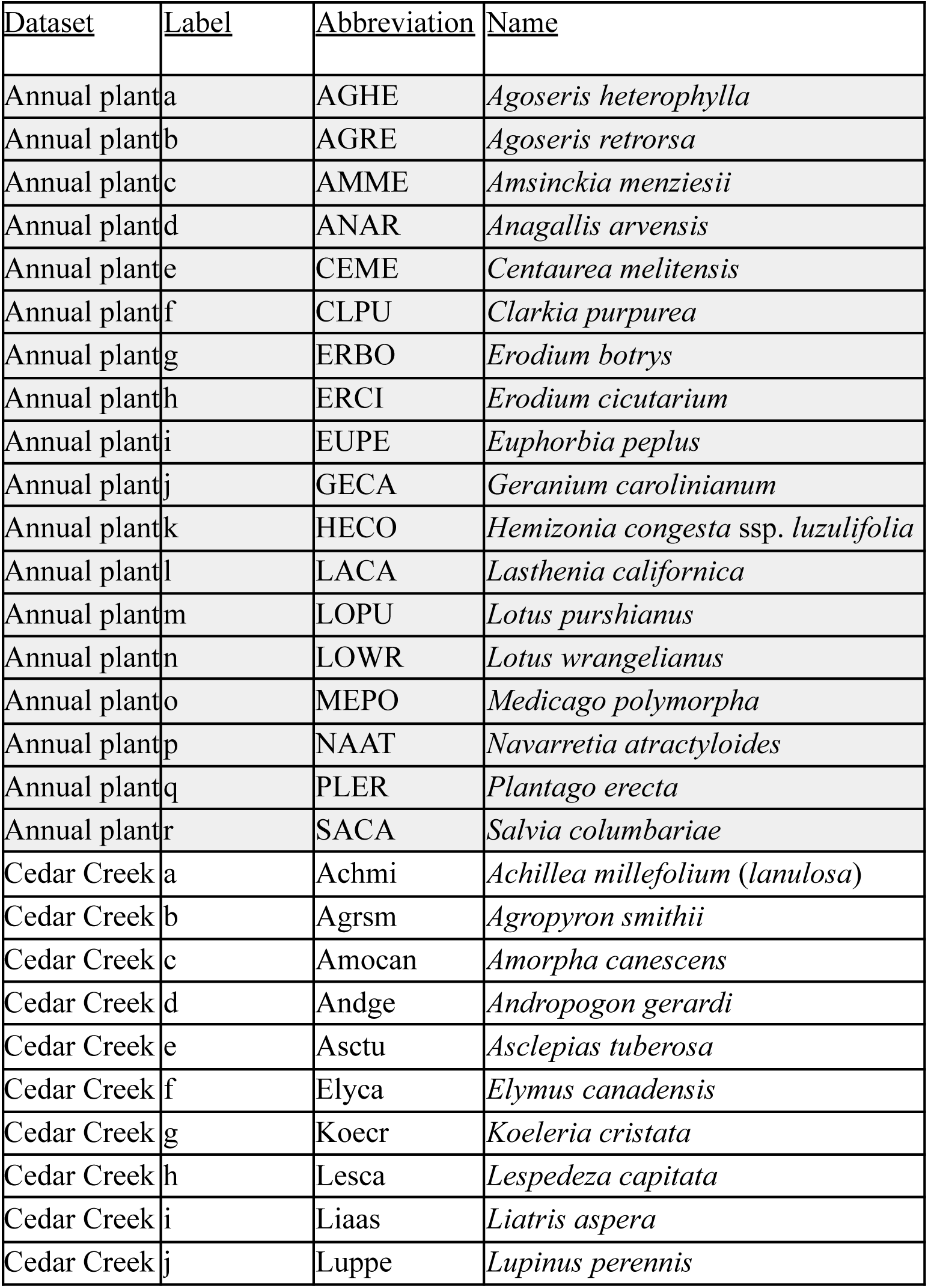
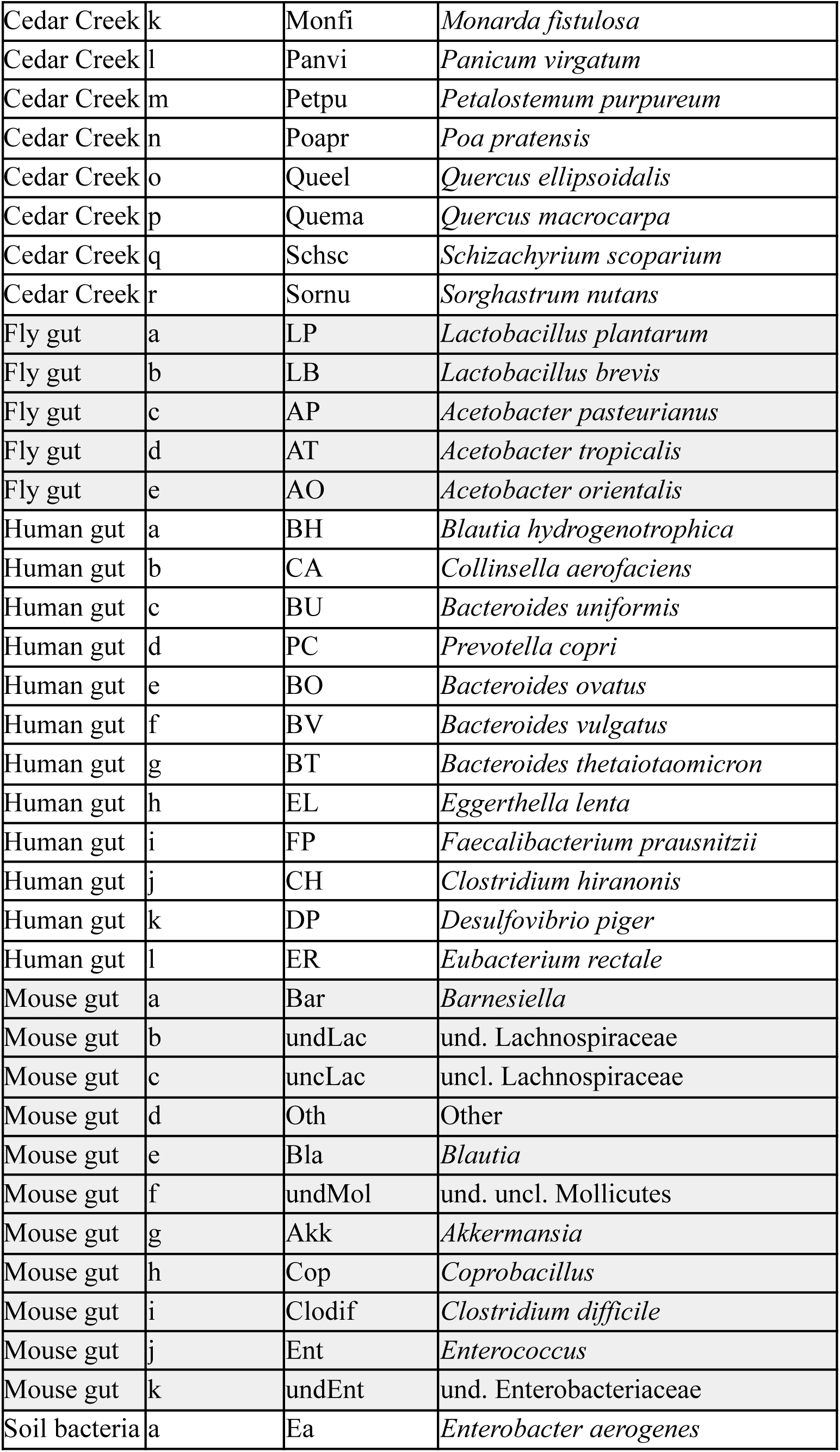
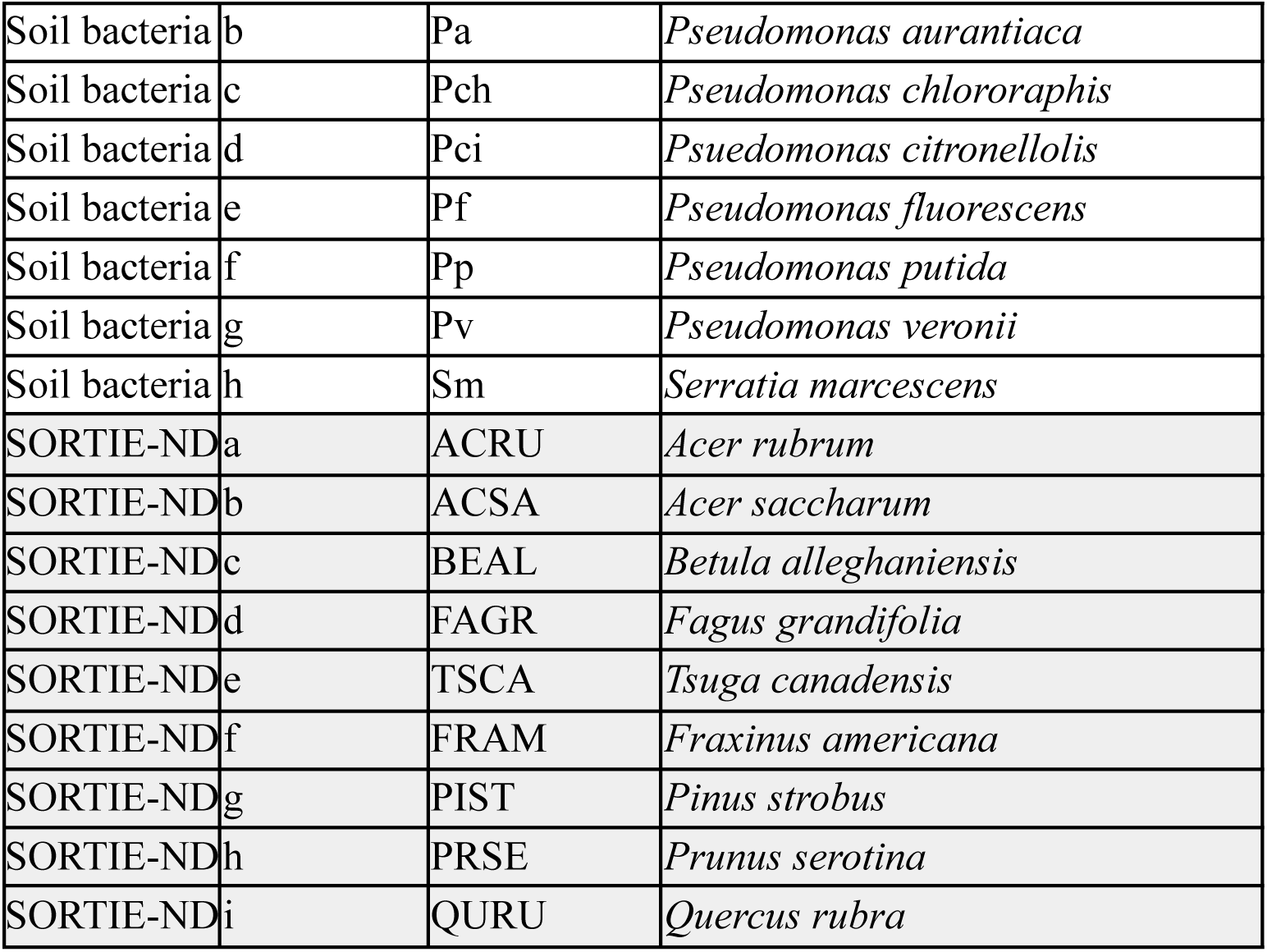
Species names for all taxa in each dataset.

**Table S2.** Scaled error for prediction, conditioned on a method of *random forest* and an experimental design of *mixed*.

| **Dataset name** | **Number of training cases** | **Scaled error (mean)** | **Scaled error (s.d.)** |
| --- | --- | --- | --- |
| Annual plant | 10 | 0.188 | 0.035206 |
| Annual plant | 14 | 0.137 | 0.051771 |
| Annual plant | 21 | 0.103 | 0.040066 |
| Annual plant | 30 | 0.08 | 0.030029 |
| Annual plant | 43 | 0.067 | 0.020783 |
| Annual plant | 62 | 0.057 | 0.015806 |
| Annual plant | 89 | 0.051 | 0.008632 |
| Annual plant | 127 | 0.048 | 0.006375 |
| Annual plant | 183 | 0.048 | 0.004666 |
| Annual plant | 264 | 0.044 | 0.005761 |
| Annual plant | 379 | 0.042 | 0.002672 |
| Annual plant | 546 | 0.043 | 0.004031 |
| Annual plant | 785 | 0.041 | 0.001772 |
| Annual plant | 1129 | 0.041 | 0.002085 |
| Annual plant | 1624 | 0.04 | 0.001111 |
| Annual plant | 2336 | 0.04 | 0.000902 |
| Annual plant | 3360 | 0.039 | 0.001127 |
| Annual plant | 4833 | 0.039 | 0.000857 |
| Annual plant | 6952 | 0.038 | 0.000657 |
| Annual plant | 10000 | 0.038 | 0.000641 |
| Cedar Creek | 10 | 0.126 | 0.007185 |
| Cedar Creek | 14 | 0.113 | 0.004621 |
| Cedar Creek | 21 | 0.105 | 0.003632 |
| Cedar Creek | 30 | 0.1 | 0.007033 |
| Cedar Creek | 43 | 0.083 | 0.002883 |
| Cedar Creek | 62 | 0.067 | 0.00401 |
| Cedar Creek | 89 | 0.045 | 0.001947 |
| Fly gut | 10 | 0.139 | 0.009458 |
| Fly gut | 14 | 0.138 | 0.007543 |
| Fly gut | 21 | 0.126 | 0.003907 |
| Fly gut | 30 | 0.127 | 0.0052 |
| Human gut | 10 | 0.134 | 0.014303 |
| Human gut | 14 | 0.111 | 0.012693 |
| Human gut | 21 | 0.103 | 0.013476 |
| Human gut | 30 | 0.084 | 0.006531 |
| Human gut | 43 | 0.067 | 0.005437 |
| Human gut | 62 | 0.053 | 0.002741 |
| Human gut | 89 | 0.043 | 0.002965 |
| Human gut | 127 | 0.035 | 0.001275 |
| Human gut | 183 | 0.029 | 0.002661 |
| Human gut | 264 | 0.025 | 0.000948 |
| Human gut | 379 | 0.02 | 0.001438 |
| Human gut | 546 | 0.018 | 0.000661 |
| Human gut | 785 | 0.016 | 0.000356 |
| Human gut | 1129 | 0.015 | 0.000273 |
| Human gut | 1624 | 0.014 | 0.000153 |
| Human gut | 2336 | 0.014 | 0.000105 |
| Human gut | 3360 | 0.014 | 0.000036 |
| Mouse gut | 10 | 0.043 | 0.00677 |
| Mouse gut | 14 | 0.033 | 0.005846 |
| Mouse gut | 21 | 0.027 | 0.002267 |
| Mouse gut | 30 | 0.024 | 0.003599 |
| Mouse gut | 43 | 0.021 | 0.001956 |
| Mouse gut | 62 | 0.018 | 0.001034 |
| Mouse gut | 89 | 0.017 | 0.001555 |
| Mouse gut | 127 | 0.017 | 0.001378 |
| Mouse gut | 183 | 0.016 | 0.001358 |
| Mouse gut | 264 | 0.015 | 0.000694 |
| Mouse gut | 379 | 0.015 | 0.000892 |
| Mouse gut | 546 | 0.015 | 0.000459 |
| Mouse gut | 785 | 0.015 | 0.000645 |
| Mouse gut | 1129 | 0.015 | 0.000207 |
| Mouse gut | 1624 | 0.015 | 0.000184 |
| SORTIE-ND | 10 | 0.101 | 0.010968 |
| SORTIE-ND | 14 | 0.092 | 0.012706 |
| SORTIE-ND | 21 | 0.085 | 0.012665 |
| SORTIE-ND | 30 | 0.071 | 0.005861 |
| SORTIE-ND | 43 | 0.063 | 0.003379 |
| SORTIE-ND | 62 | 0.056 | 0.001947 |
| SORTIE-ND | 89 | 0.052 | 0.001347 |
| SORTIE-ND | 127 | 0.051 | 0.001218 |
| SORTIE-ND | 183 | 0.048 | 0.000681 |
| SORTIE-ND | 264 | 0.045 | 0.000476 |
| SORTIE-ND | 379 | 0.042 | 0.000352 |
| Soil bacteria | 10 | 0.136 | 0.006685 |
| Soil bacteria | 14 | 0.125 | 0.007807 |
| Soil bacteria | 21 | 0.106 | 0.011251 |
| Soil bacteria | 30 | 0.084 | 0.009954 |
| Soil bacteria | 43 | 0.065 | 0.003127 |
| Soil bacteria | 62 | 0.049 | 0.00419 |
| Soil bacteria | 89 | 0.034 | 0.002287 |

**Table S3.** Error for prioritization, conditioned on a method of *random forest* and an experimental design of *mixed*.

| **Task** | **Dataset name** | **Number of training cases** | **True negative rate (mean)** | **True negative rate (s.d.)** | **True positive rate (mean)** | **True positive rate (s.d.)** |
| --- | --- | --- | --- | --- | --- | --- |
| abundance | Annual plant | 89 | 0.000031 | 0.000096 | 0.99 | 0.01719 |
| abundance | Human gut | 89 | 0.472195 | 0.303772 | 0.99 | 0.00522 |
| abundance | Mouse gut | 89 | 0 | 0 | 1 | 0 |
| abundance | SORTIE-N D | 89 | 0.020968 | 0.017086 | 0.99 | 0.01031 |
| abundance | Annual plant | 264 | 0 | 0 | 1 | 0.00024 |
| abundance | Human gut | 264 | 0.462927 | 0.306334 | 0.99 | 0.00414 |
| abundance | Mouse gut | 264 | 0 | 0 | 1 | 0 |
| abundance | SORTIE-N D | 264 | 0.032258 | 0.026339 | 0.98 | 0.00581 |
| shannons_h | Annual plant | 89 | 0.247177 | 0.152217 | 0.97 | 0.02131 |
| shannons_h | Human gut | 89 | 0.598206 | 0.197346 | 0.98 | 0.01353 |
| shannons_h | Mouse gut | 89 | 0.287778 | 0.111167 | 1 | 0.00174 |
| shannons_h | SORTIE-N D | 89 | 0.039706 | 0.028625 | 0.99 | 0.00339 |
| shannons_h | Annual plant | 264 | 0.71644 | 0.112456 | 0.95 | 0.00499 |
| shannons_h | Human gut | 264 | 0.653812 | 0.17978 | 0.98 | 0.01621 |
| shannons_h | Mouse gut | 264 | 0.373333 | 0.151009 | 0.99 | 0.00442 |
| shannons_h | SORTIE-N D | 264 | 0.108824 | 0.030376 | 0.99 | 0.00529 |
| removal | Annual plant | 89 | 0.85757 | 0.219518 | 0.97 | 0.05003 |
| removal | Human gut | 89 | 0.902874 | 0.168068 | 0.91 | 0.06902 |
| removal | Mouse gut | 89 | 0.776574 | 0.284374 | 0.94 | 0.04922 |
| removal | SORTIE-N<br>D | 89 | 0.78989 | 0.271564 | 0.94 | 0.06289 |
| removal | Annual<br>plant | 264 | 0.863898 | 0.198121 | 0.99 | 0.01778 |
| removal | Human gut | 264 | 0.912263 | 0.098506 | 0.96 | 0.03929 |
| removal | Mouse gut | 264 | 0.777913 | 0.284855 | 0.98 | 0.02 |
| removal | SORTIE-N<br>D | 264 | 0.747148 | 0.293548 | 0.97 | 0.02973 |

## Text S1

Datasets and pre-processing steps.

### ‘Annual plant’ dataset

We obtained data from a field-parameterized plant competition model, which describes the dynamics of annual plants with seed banks (Chesson 1990; Levine & HilleRisLambers 2009). This model is more complex than the generalized Lotka-Volterra, as it includes population stage structure and nonlinear competition. Data came from 18 California annual plants (Godoy *et al*. 2014). We modified the model reported in (Godoy *et al*. 2014) to include multi-species competition, following (Godoy *et al*. 2017). The modified discrete-time model describes the abundance of seeds of species *i* at time *t+1* as:

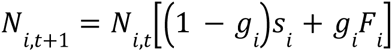

where:

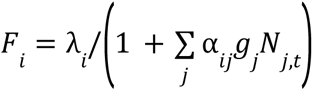

The modification is the inclusion in the denominator of *F_i_* of a sum over all species, rather than the sum over only two focal species. Here, λ*_i_* is the per germinant fecundity of species *i*,*g_i_* is the germination rate of species *i*, *s_i_* is the annual survival rate of ungerminated seed in the soil of species *i*, *F_i_* is the number of viable seeds produced per germinated individual of species *i*, and α*_ij_* is the per capita effect of species *j* on species *i*. 66/234 values of α*_ij_* which were missing from the dataset were replaced with the mean value in the dataset per (Donders *et al*. 2006).

For each of the communities possible among the species pool, we initialized all species present in the experiment to *N_i_* (*t* = 0) = 1, and ran for 1000 generations (long enough to reach equilibrium). Richness and composition were calculated by flagging species with *N_i_* (*t* = 1000)≥0. 01.

### ‘Cedar Creek’ dataset

We obtained data from the Cedar Creek Biodiversity II ‘e120’ experiment. This dataset describes annual aboveground biomass estimates (from 1994 to 2018) of 154 experimentally assembled plant communities of varying composition (Tilman *et al*. 2001). For each plot, we set the experimental conditions (X) to whether the plot contained each of *n*=18 species (16 intentionally planted, plus 2 volunteer species). We then set the final abundance to each species’ biomass in each plot in 2018. Richness and composition were calculated by flagging species with *N_i_* (*t* = 2018) > 0. This approach conflates biomass with abundance and does not account for biomass from other non-focal species that colonized each plot by 2018 (e.g. numerous weeds), but is a reasonable choice given the limitations of the data.

### ‘Fly gut’ dataset

We obtained data for germ-free fruit flies experimentally inoculated with each possible combination of core species of fly gut bacteria at 3-day intervals, from (Gould *et al*. 2018). For each treatment, we set the experimental conditions (X) to whether the fly had been inoculated with each of *n*=5 bacterial taxa. We then set the final abundances to the number of colony forming units of each taxon after 10 days of experimental treatments. A total of *q=*48 replicate flies were used per treatment. Richness and composition were calculated by flagging species with *N_i_* (*t* = 10) > 0.

### ‘Human gut’ dataset

The generalized Lotka-Volterra model was used to simulate outcomes, based on the equation:

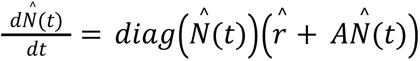

Here 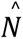 is a vector of abundances among species in the regional pool, 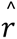 is the vector of density-independent growth rates, and *A* is the matrix of interaction coefficients, with entry *A_ij_*representing the change in species *i*’s per-capita growth rate for a unit change in the density of species *j*.

For each of the communities possible among the species pool, and for a given set of *A* and *r* parameters, we analyzed the model over the reduced dimensionality corresponding to the number of species introduced in the local community. We initialized all species present in the experiment to *N_i_*(*t* = 0) = 1, then solved the differential equation up to *t*=1000 using the *ode* function in the *deSolve* package in R. Richness and composition were calculated by flagging species with 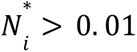.

We parameterized the model for a *n*=11 mouse gut microbial community including the pathogen *Clostridium difficile* (Stein *et al*. 2013) based on (Buffie *et al*. 2012).

### ‘Mouse gut’ dataset

We followed the steps outlined for the ‘human gut’ dataset but using *A* and *r* parameters for a *n*=12 synthetic human gut microbial community (Venturelli *et al*. 2018).

### ‘Soil bacteria’ dataset

We obtained data from experimental assembly of soil bacterial communities from (Friedman *et al*. 2017). Communities were assembled at varying densities in microplate microcosms each comprising five cycles each comprising 48 hours of growth, followed by a 1500-fold dilution into fresh media. Data include species grown alone, in pairs, in triplets, in single-species drop-outs, and all together. Experiments were replicated from 2 to 14 times. We set the abundance of each species to its optical density after this growth period. Richness and composition were calculated by flagging species with *N_i_* (*t* = 240) > 0.

### ‘SORTIE-ND’ dataset

We used the SORTIE-ND (version 7.0.5) model of forest dynamics, which is an individual-based forest simulator that includes demography and life history stage transitions, light competition, spatially explicit dispersal, and other processes (Pacala *et al*. 1993, 1996). This model was chosen for its high complexity and stochasticity.

We obtained a parameterization of the model for *n*=9 hardwood species in eastern North America at 42°N latitude (‘GMD’, available by download from http://sortie-nd.org/software/7_05/sample_par_file.zip). We modified this file to change the plot size to 100 x 100 m, to run for 1000 years (200 5-year time steps) with no external disturbances, and set the parameters for Weibull seed rain and Weibull seed beta to species-specific values reported at http://sortie-nd.org/software/sample_par_file.html), as the default parameter file erroneously includes blank values (personal communication, L. Murphy and C. Canham, 2 Sept. 2021). The simulation does not come to equilibrium but rather includes fluctuations in abundance, due to the effects of light-based competition and no self-thinning in the understorey. Some stochastic extinctions also occur.

For each local community that could be assembled from this species pool, we then ran simulations, initializing all species to an initial density of either 0 or 25 saplings ha^-1^ (default initial values) and running *q*=3 replicates per initial condition. We determined abundance as the absolute density of adults at *t*=1000. Richness and composition were calculated by flagging species with adult densities of *N_*i*_*(*t* = 1000) > 1.

---

**Figure S1.**
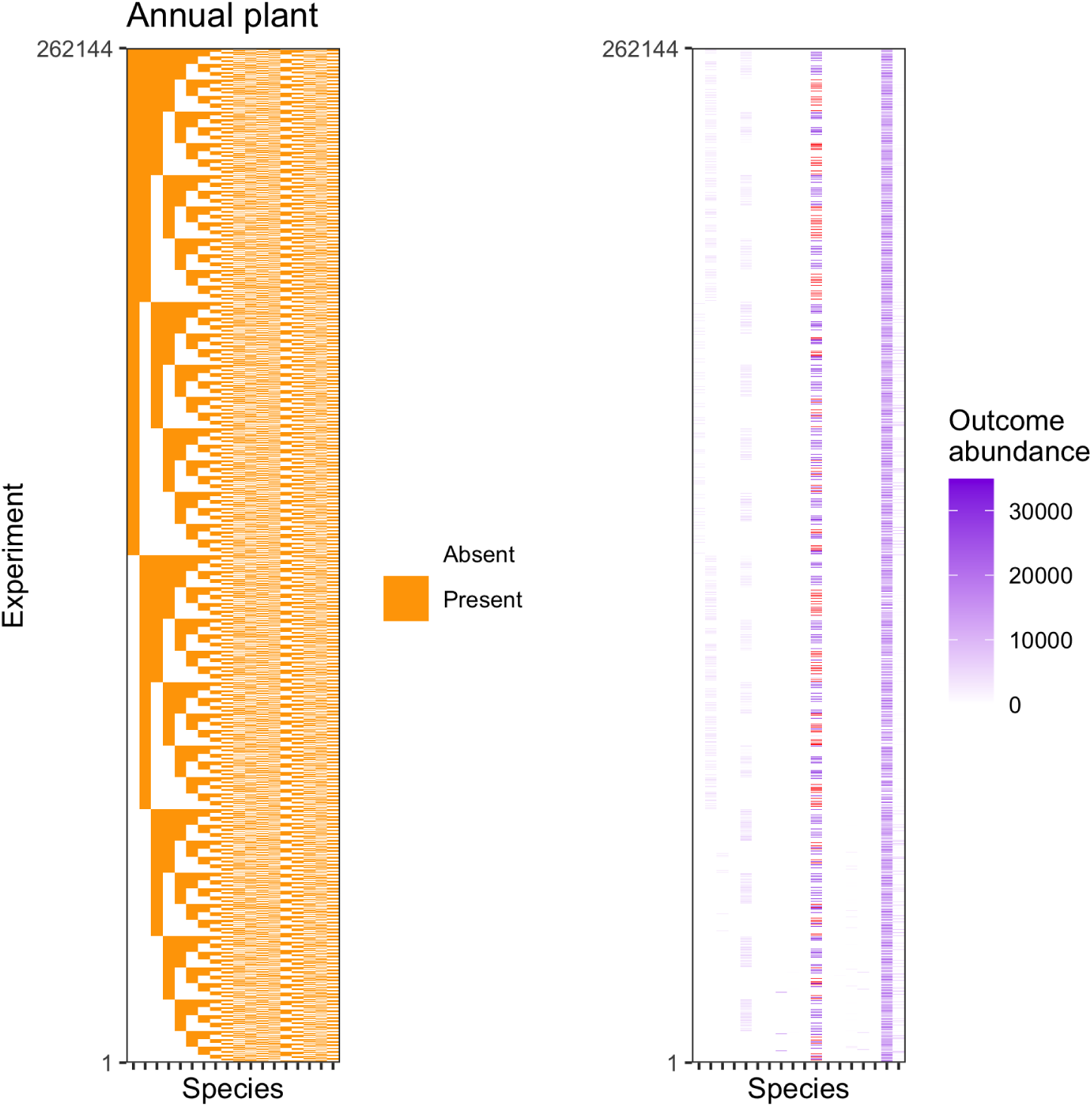
Visualization of experimental conditions and abundance outcomes for the annual plant dataset. Panels show **(a)** initial species presence/absence data for each experiment and **(b)** outcomes. Quantile clipped values are colored red.

**Figure S2.**
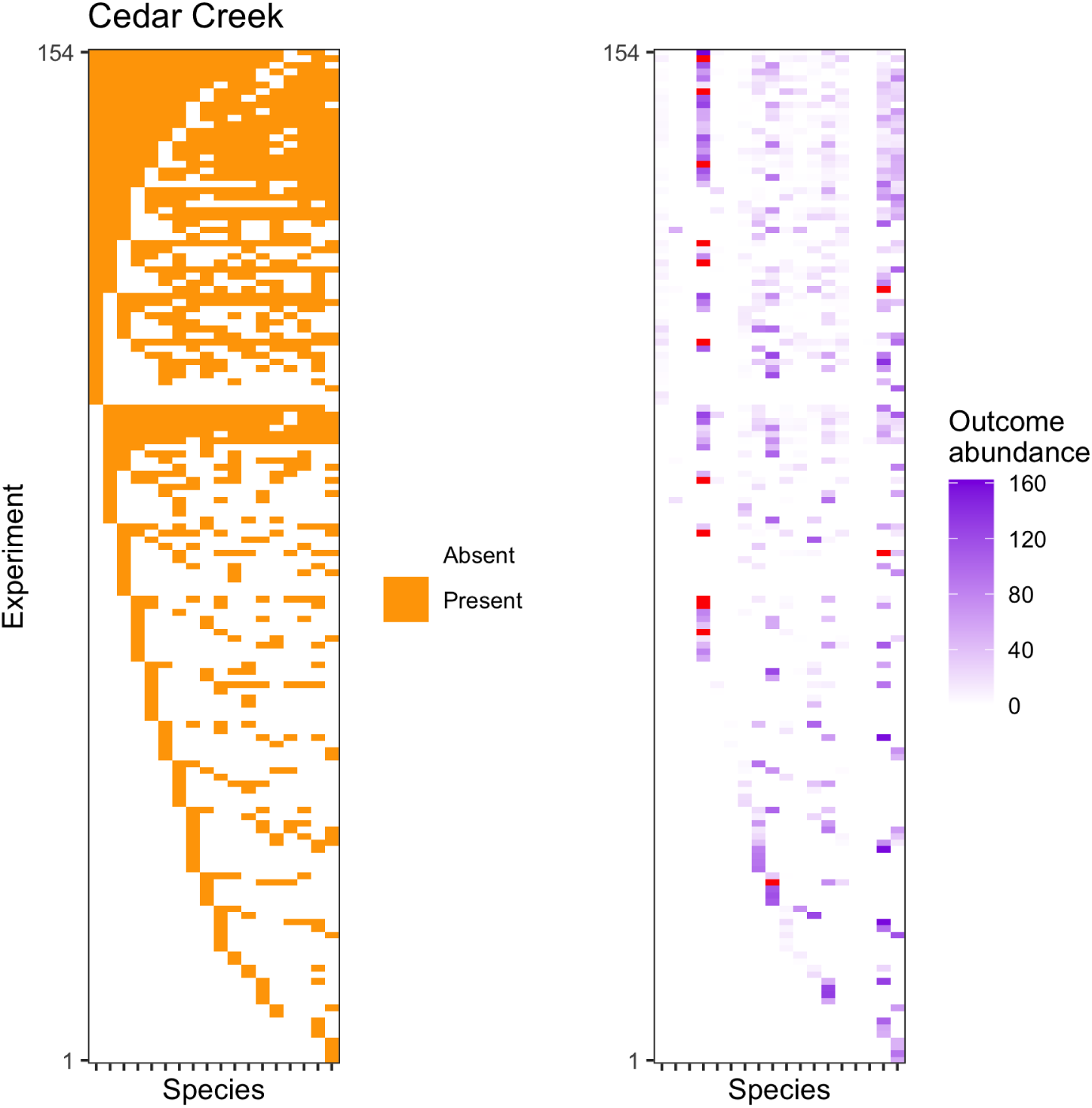
Visualization of experimental conditions and abundance outcomes for the Cedar Creek dataset. Panels show **(a)** initial species presence/absence data for each experiment and **(b)** outcomes. Quantile clipped values are colored red.

**Figure S3.**
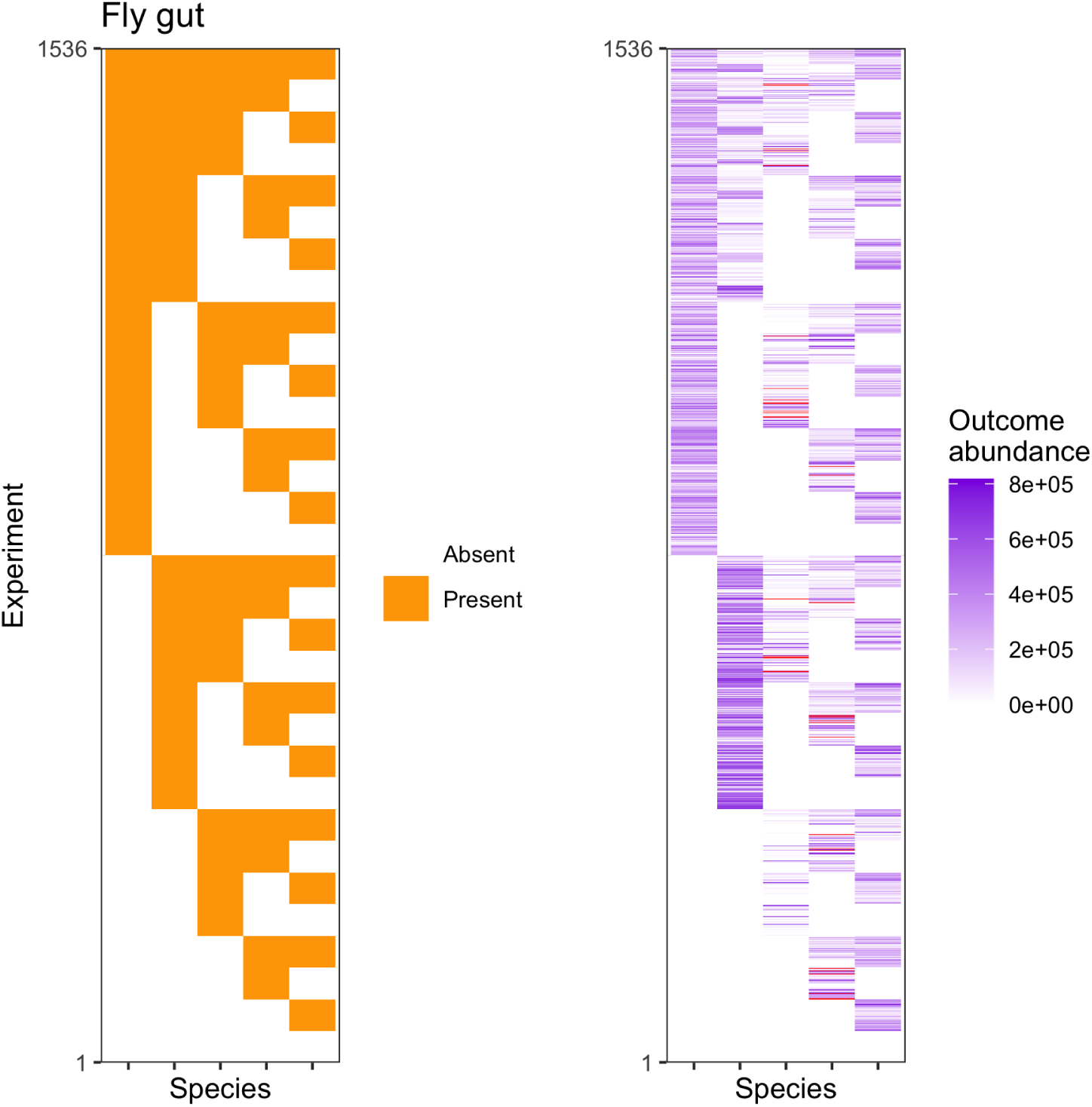
Visualization of experimental conditions and abundance outcomes for the fly gut dataset. Panels show **(a)** initial species presence/absence data for each experiment and **(b)** outcomes. Quantile clipped values are colored red.

**Figure S4.**
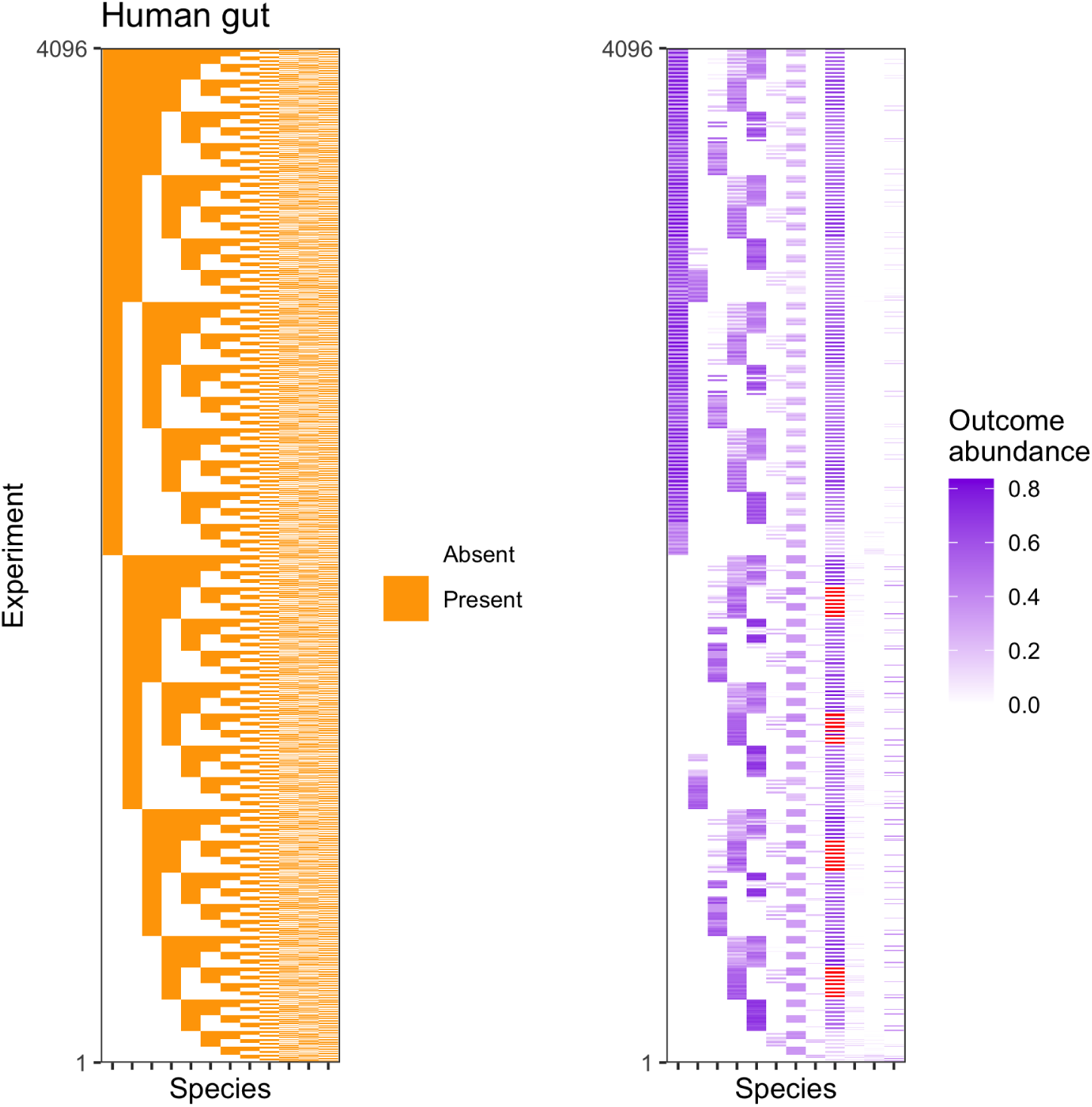
Visualization of experimental conditions and abundance outcomes for the human gut dataset. Panels show **(a)** initial species presence/absence data for each experiment and **(b)** outcomes. Quantile clipped values are colored red.

**Figure S5.**
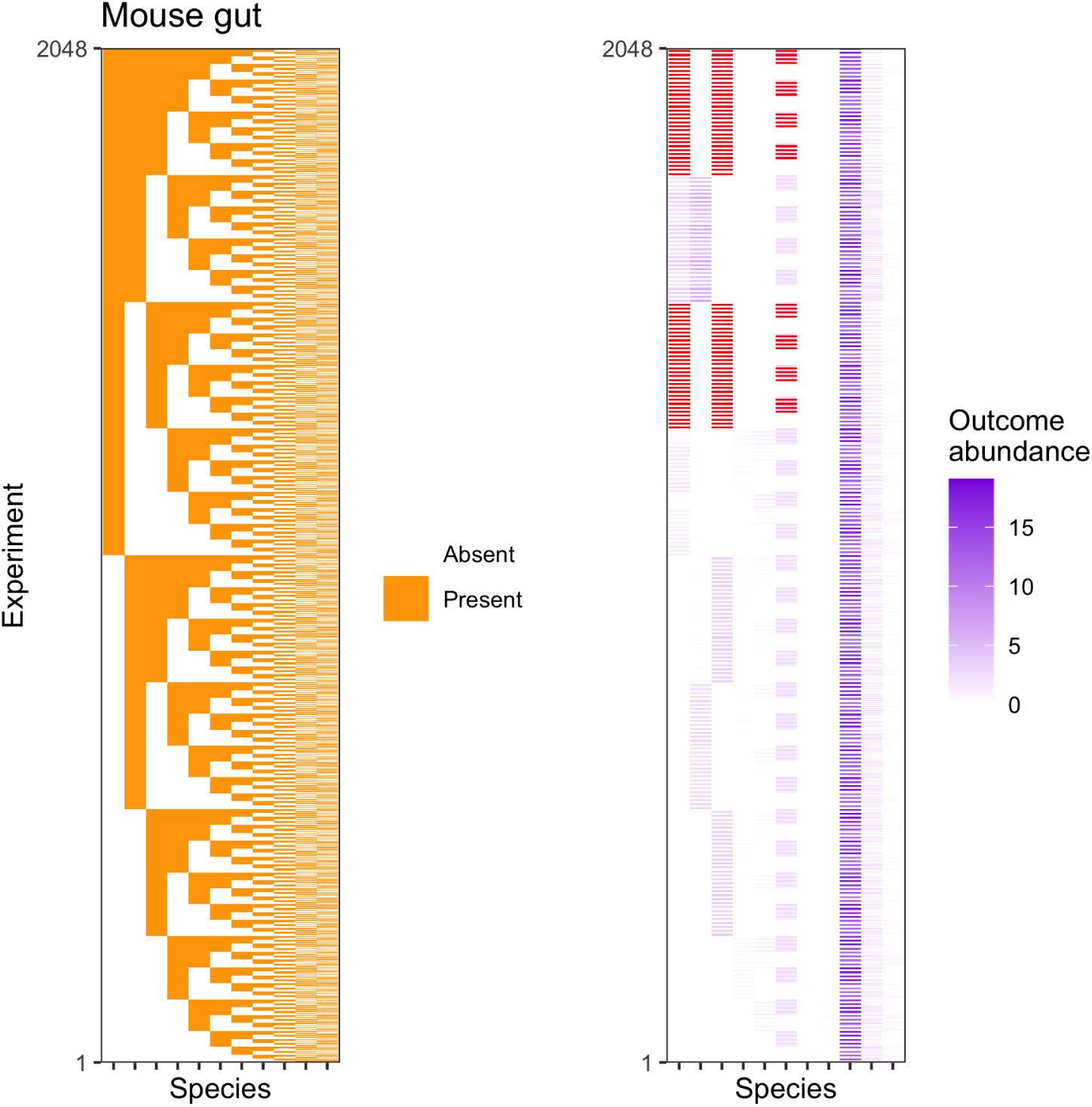
Visualization of experimental conditions and abundance outcomes for the mouse gut dataset. Panels show **(a)** initial species presence/absence data for each experiment and **(b)** outcomes. Quantile clipped values are colored red.

**Figure S6.**
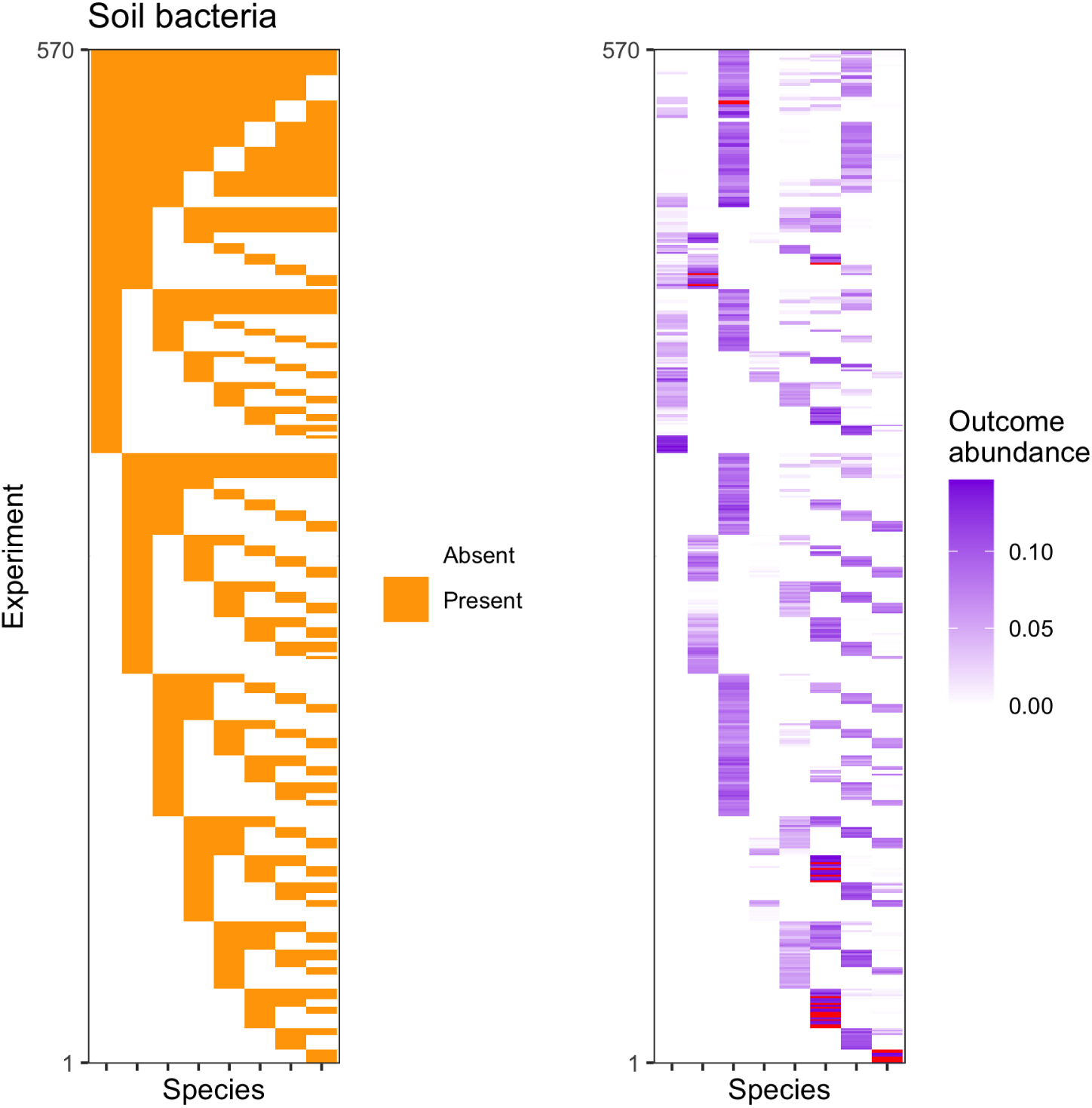
Visualization of experimental conditions and abundance outcomes for the soil bacteria dataset. Panels show **(a)** initial species presence/absence data for each experiment and **(b)** outcomes. Quantile clipped values are colored red.

**Figure S7.**
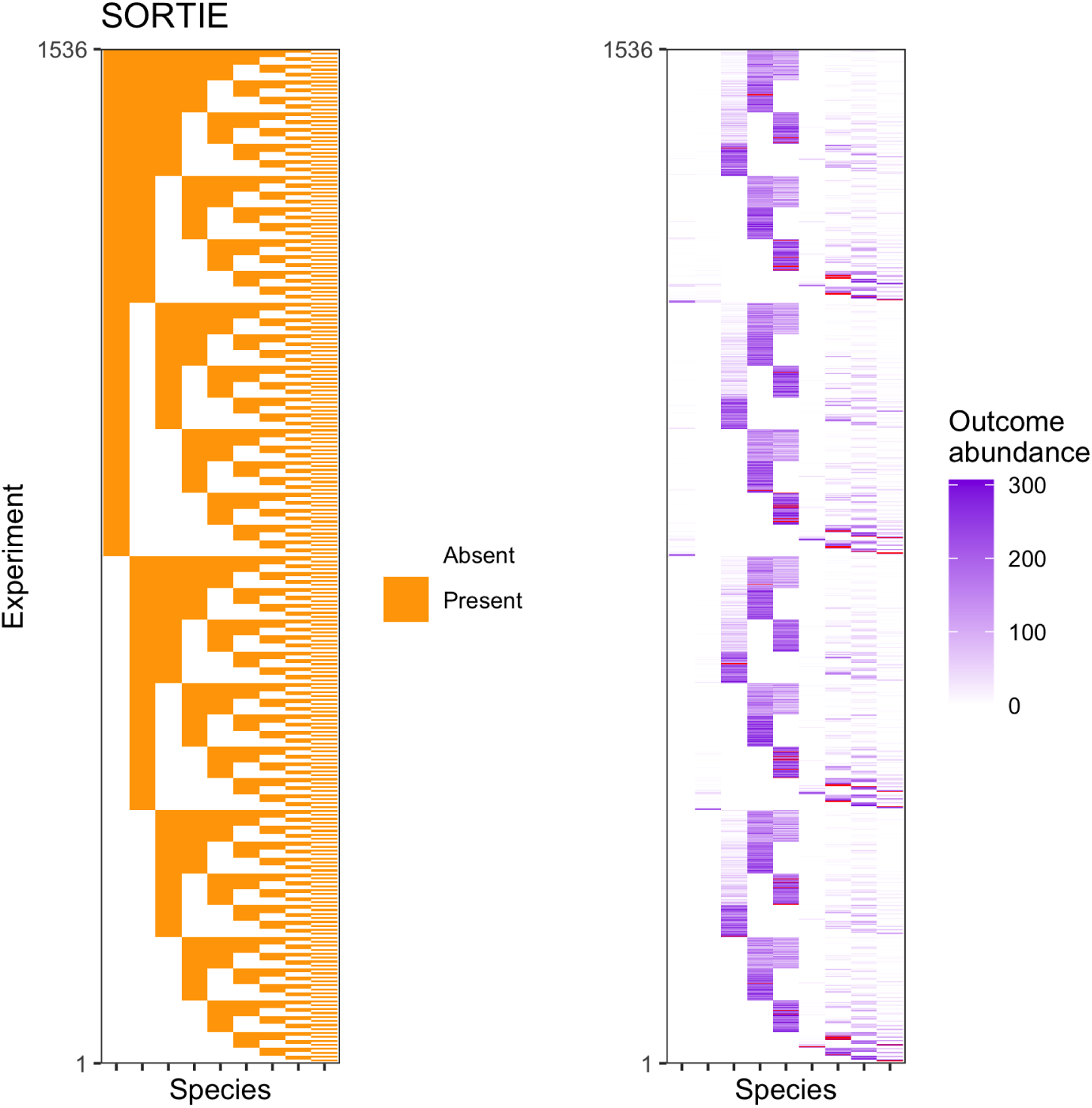
Visualization of experimental conditions and abundance outcomes for the SORTIE dataset. Panels show **(a)** initial species presence/absence data for each experiment and **(b)** outcomes. Quantile clipped values are colored red.

**Figure S8.**
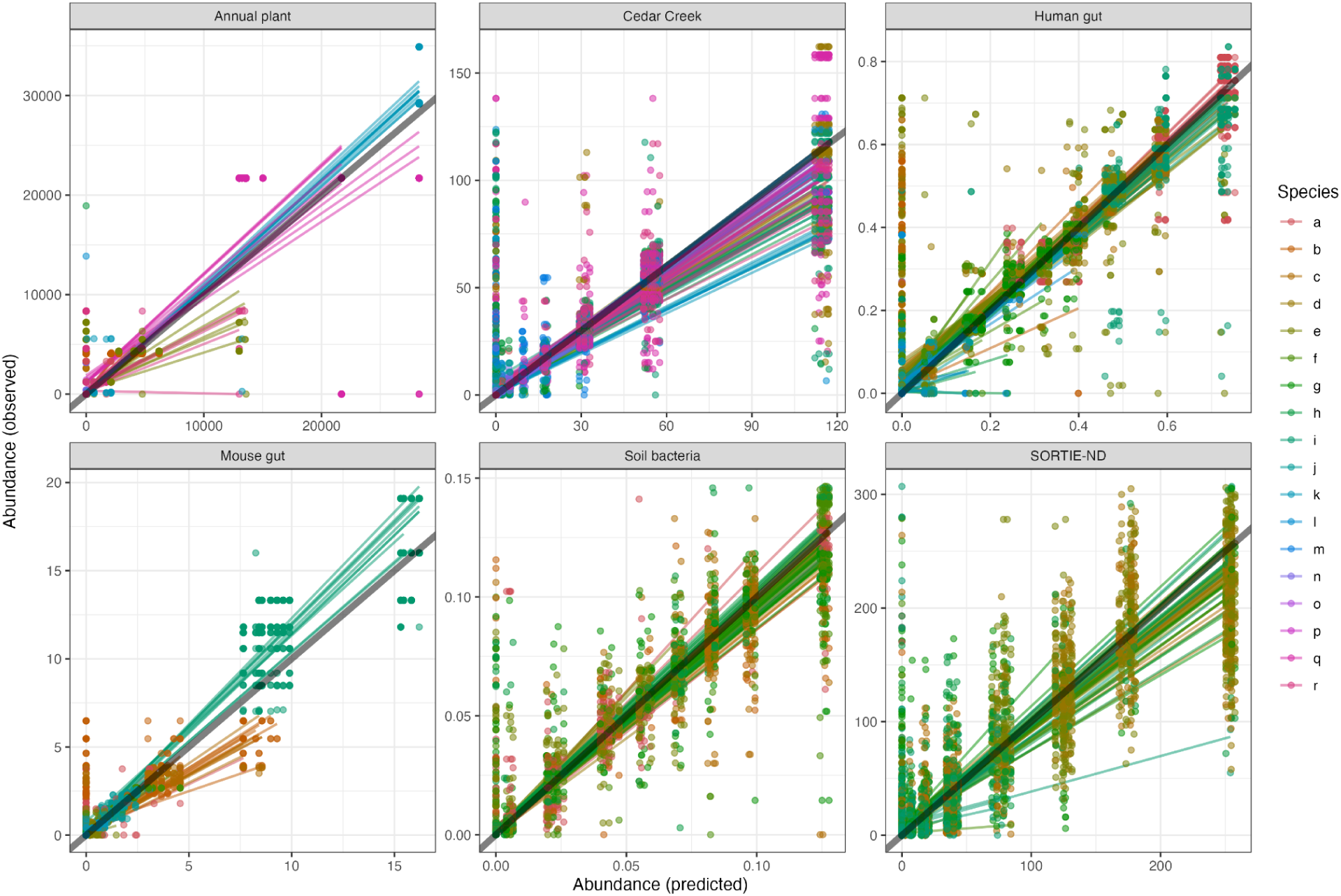
Observed vs. predicted abundance values for all species and all training replicates. Individual predictions are shown as dots; lines are drawn for each species and replicate sample dataset combination, and reflect a regression for all test-set experiments of this combination. The 1:1 line is shown in transparent black. Predictions are for a random forest method, a mixed richness experimental design, and 89 training experiments. Species names for each alphabetical species code are in Table S1.

**Figure S9.**
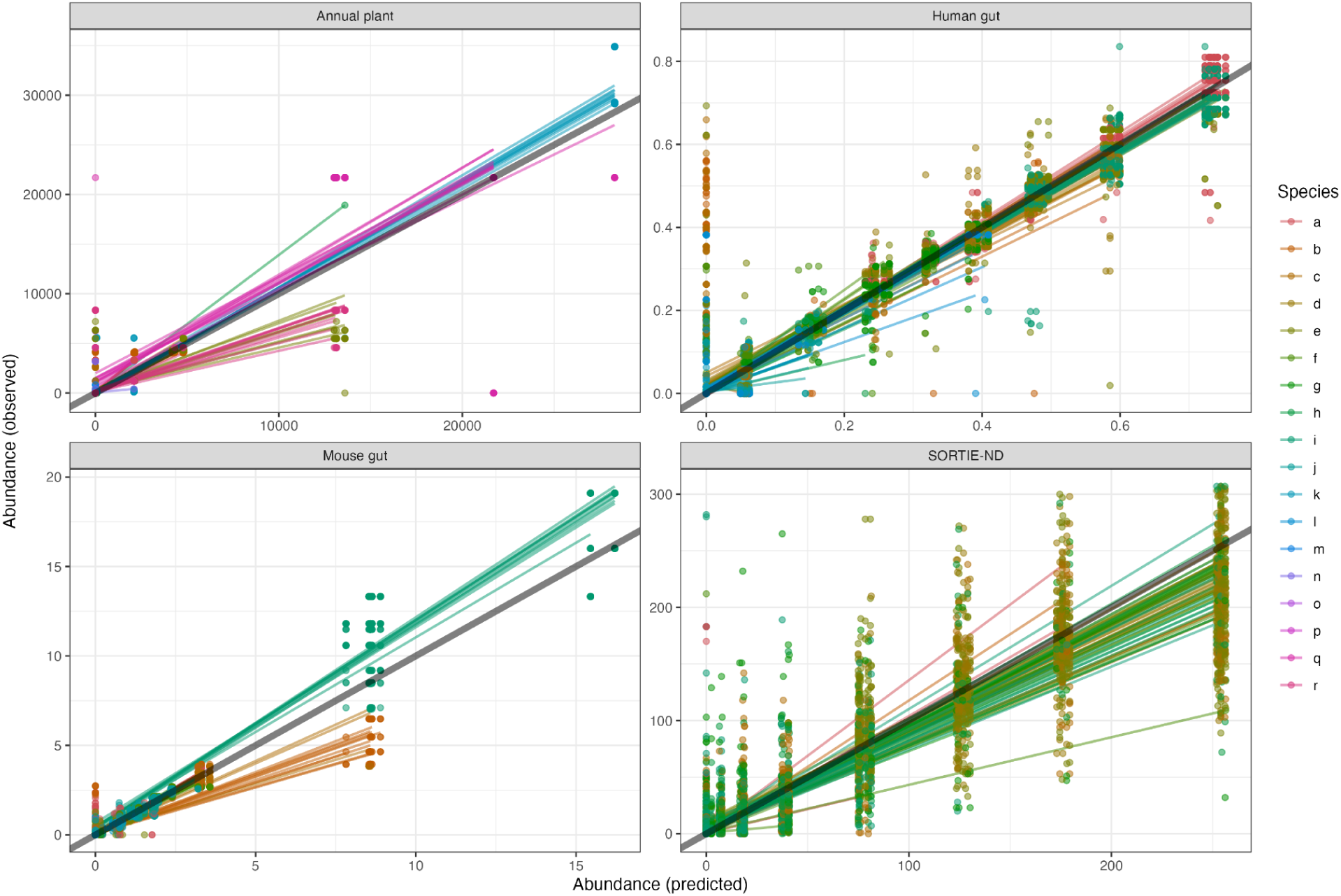
Observed vs. predicted abundance values for all species and all sampled training datasets. Individual predictions are shown as dots; lines are drawn for each species and replicate sample dataset combination, and reflect a regression for all test-set experiments of this combination. The 1:1 line is shown in transparent black. Predictions are for a random forest method, a mixed richness experimental design, and 264 training experiments. Species names for each alphabetical species code are in **Table S1**. Some datasets did not have enough training experiments at this sample size to be visualized.

**Figure S10.**
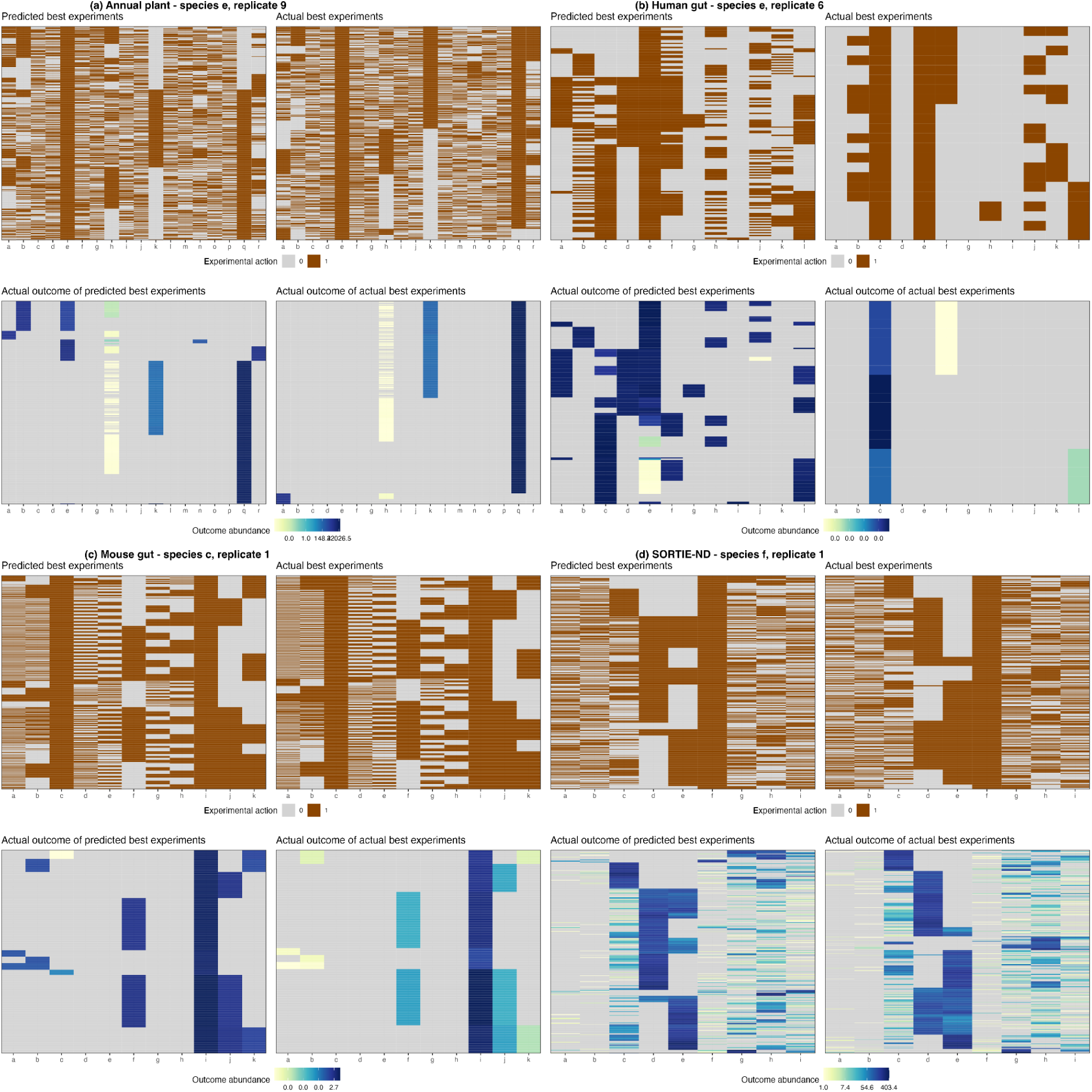
Structure of prioritization errors for the removal task for the **(a)** annual plant, **(b)** human gut, **(c)** mouse gut and **(d)** SORTIE-ND datasets. Within each panel, left columns indicate prioritizations for a random forest method, a mixed richness experimental design, and 89 training experiments and right columns indicate actual best experiments. Top panels indicate experiments as rows, while bottom panels indicate outcomes as rows. The ordering of rows in top and bottom panels is the same and is based on hierarchical clustering of the outcomes. In panel a, a random sample of 500 experiments is shown for clearer visualization. The replicate and species combination with highest true positive rate has been chosen for this visualization. Species names for each alphabetical species code are in **Table S1**.

**Figure S11.**
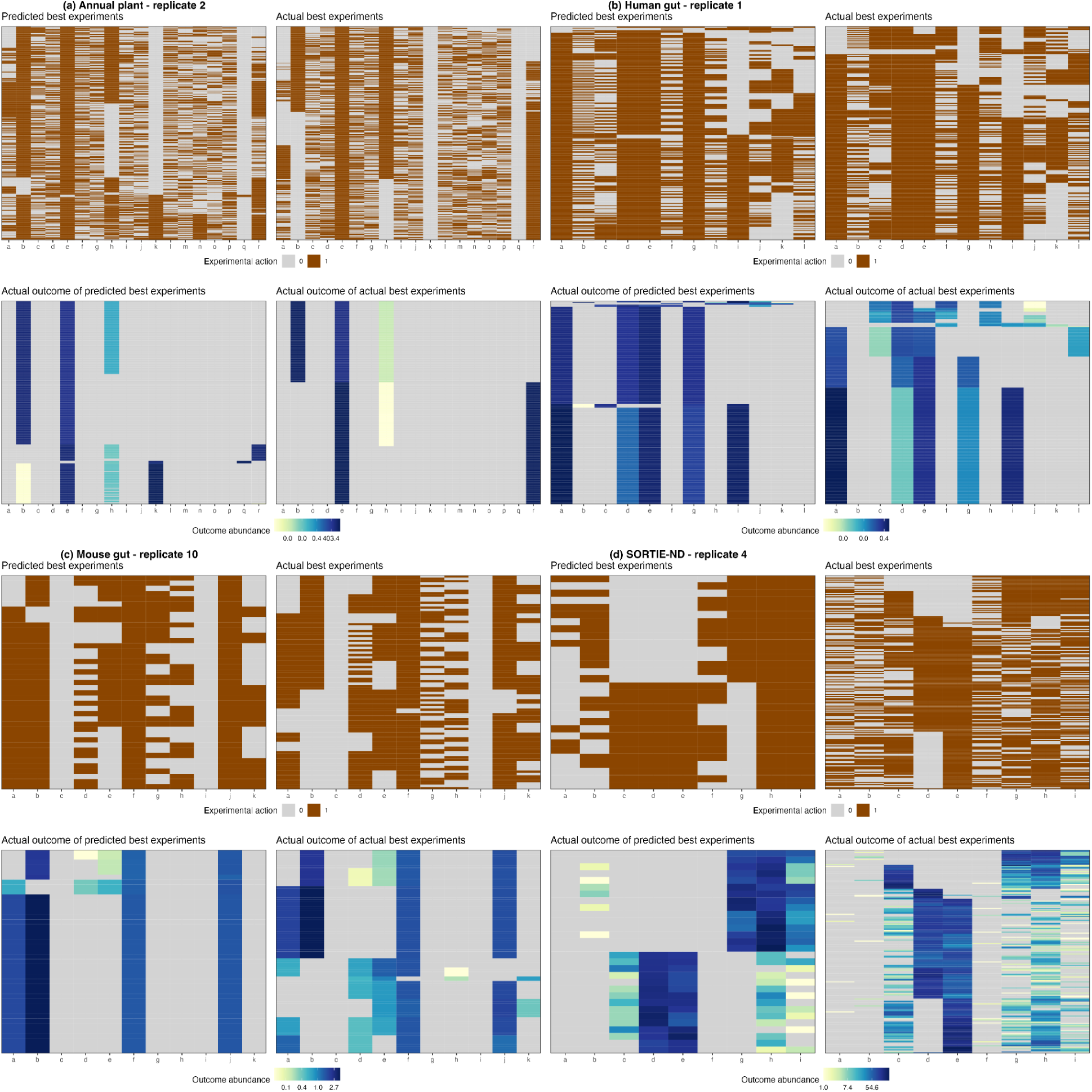
Structure of prioritization errors for the maximizing Shannon’s H task for the **(a)** annual plant, **(b)** human gut, **(c)** mouse gut and **(d)** SORTIE-ND datasets. Within each panel, left columns indicate prioritizations for a random forest method, a mixed richness experimental design, and 89 training experiments and right columns indicate actual best experiments. Top panels indicate experiments as rows, while bottom panels indicate outcomes as rows. The ordering of rows in top and bottom panels is the same and is based on hierarchical clustering of the outcomes. In panel a, a random sample of 500 experiments is shown for clearer visualization. The replicate with highest true positive rate has been chosen for this visualization. Species names for each alphabetical species code are in **Table S1**.

**Figure S12.**
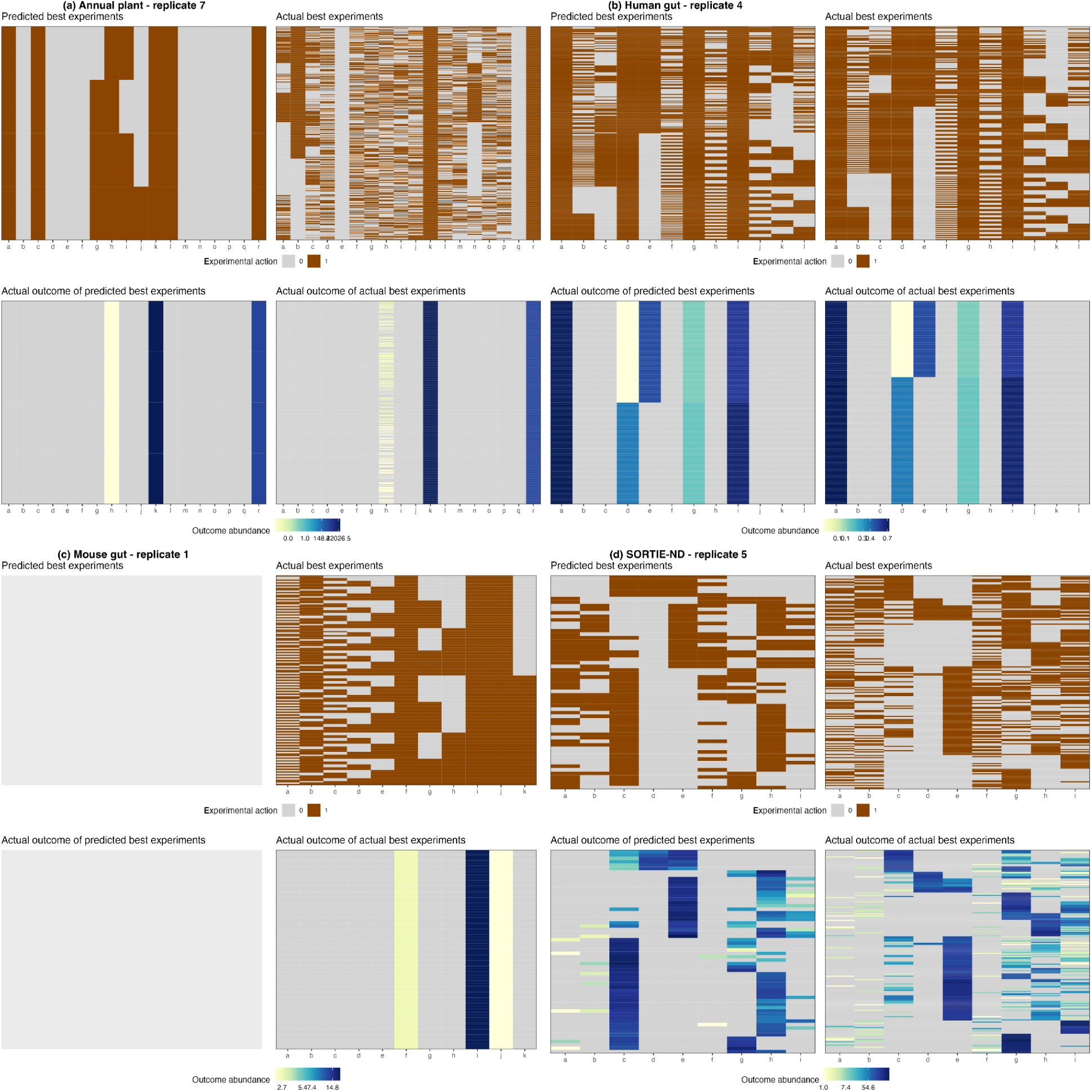
Structure of prioritization errors for the maximizing total abundance task for the **(a)** annual plant, **(b)** human gut, **(c)** mouse gut and **(d)** SORTIE-ND datasets. Within each panel, left columns indicate prioritizations for a random forest method, a mixed richness experimental design, and 89 training experiments and right columns indicate actual best experiments. Top panels indicate experiments as rows, while bottom panels indicate outcomes as rows. The ordering of rows in top and bottom panels is the same and is based on hierarchical clustering of the outcomes. In panel a, a random sample of 500 experiments is shown for clearer visualization. The replicate with highest true positive rate has been chosen for this visualization. Species names for each alphabetical species code are in **Table S1**. Blanks are shown in panel c left column due to the failure of the prioritization to identify any viable predictions.

**Figure S13.**
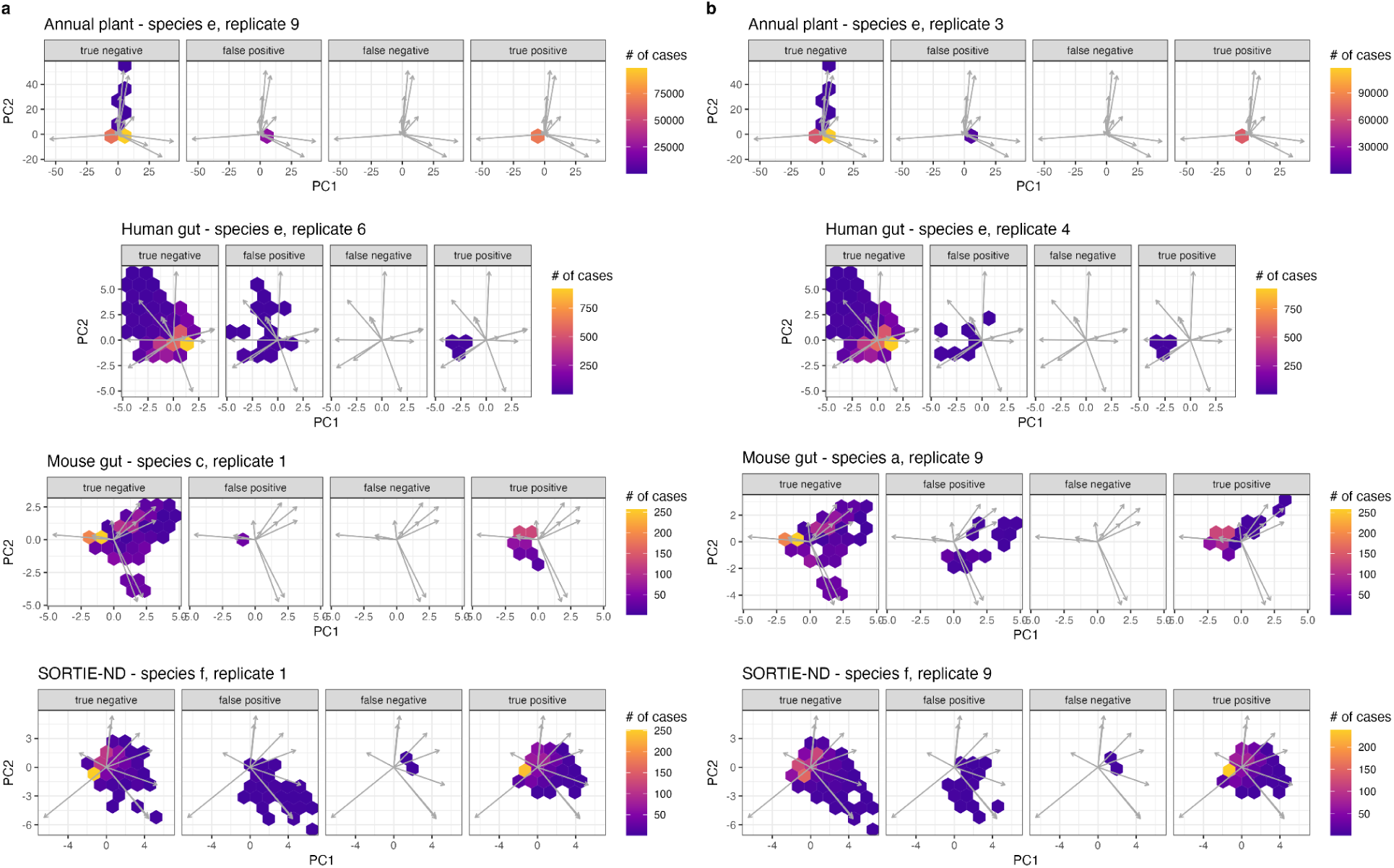
Structure of error types in abundance outcome space for the removal prioritization task for each dataset. Prioritizations are for a random forest method, a mixed richness experimental design, and **(a)** 89 or **(b)** 264 training experiments. In each panel, arrows show variable loadings of a principal component analysis in outcome abundance space (*x*^1/4^ transformed to reduce outlier effects); hexagon colors indicate numbers of outcomes that fall within each bin. Outcomes are grouped by whether the experiment yielding them is a true positive, true negative, false positive, or false negative with respect to the prioritization task. Species names for each alphabetical species code are in **Table S1**.

**Figure S14.**
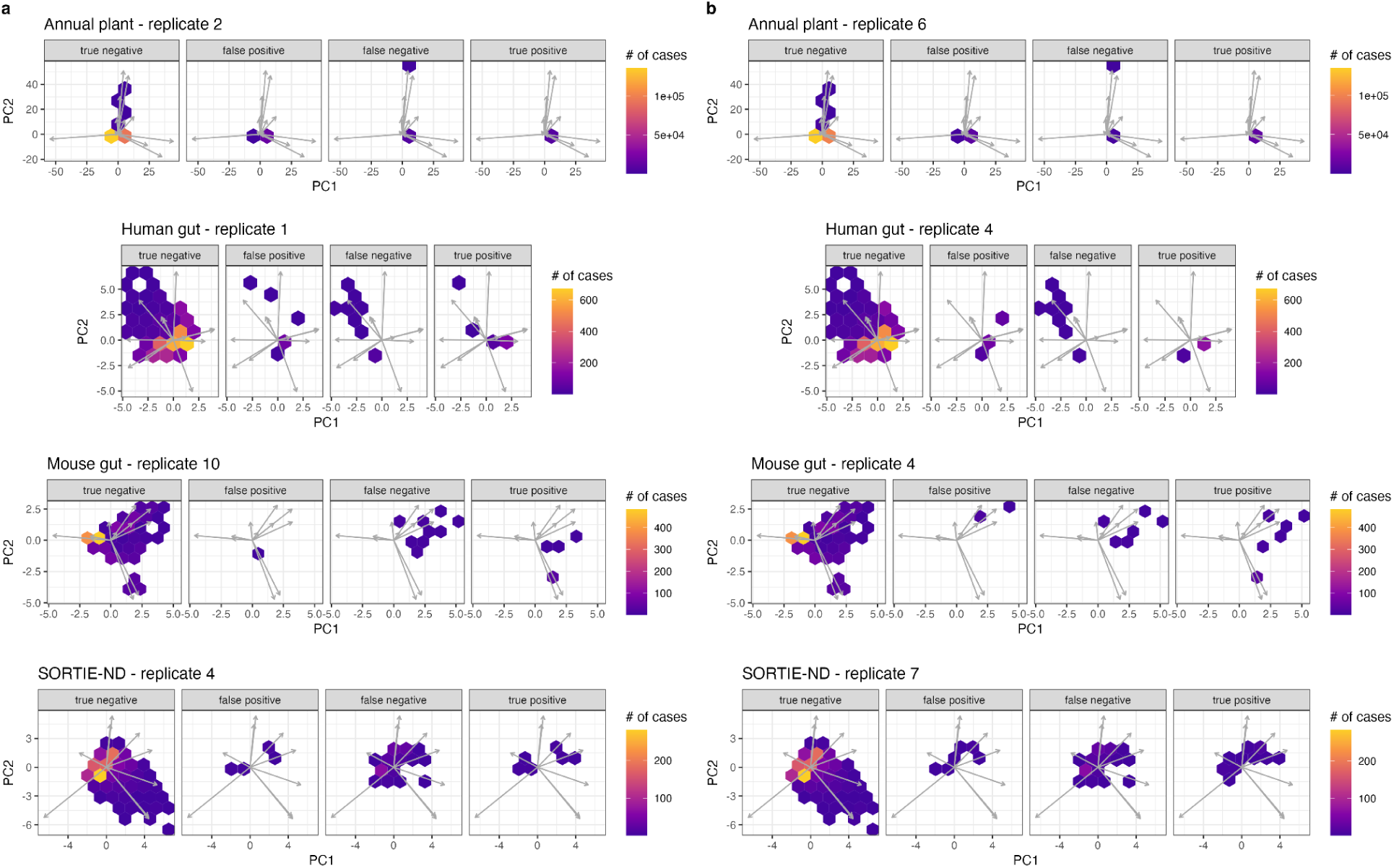
Structure of error types in abundance outcome space for the maximizing Shannon’s H task for each dataset. Prioritizations are for a random forest method, a mixed richness experimental design, and **(a)** 89 or **(b)** 264 training experiments. In each panel, arrows show variable loadings of a principal component analysis in outcome abundance space (*x*^1/4^ transformed to reduce outlier effects); hexagon colors indicate numbers of outcomes that fall within each bin. Outcomes are grouped by whether the experiment yielding them is a true positive, true negative, false positive, or false negative with respect to the prioritization task.

**Figure S15.**
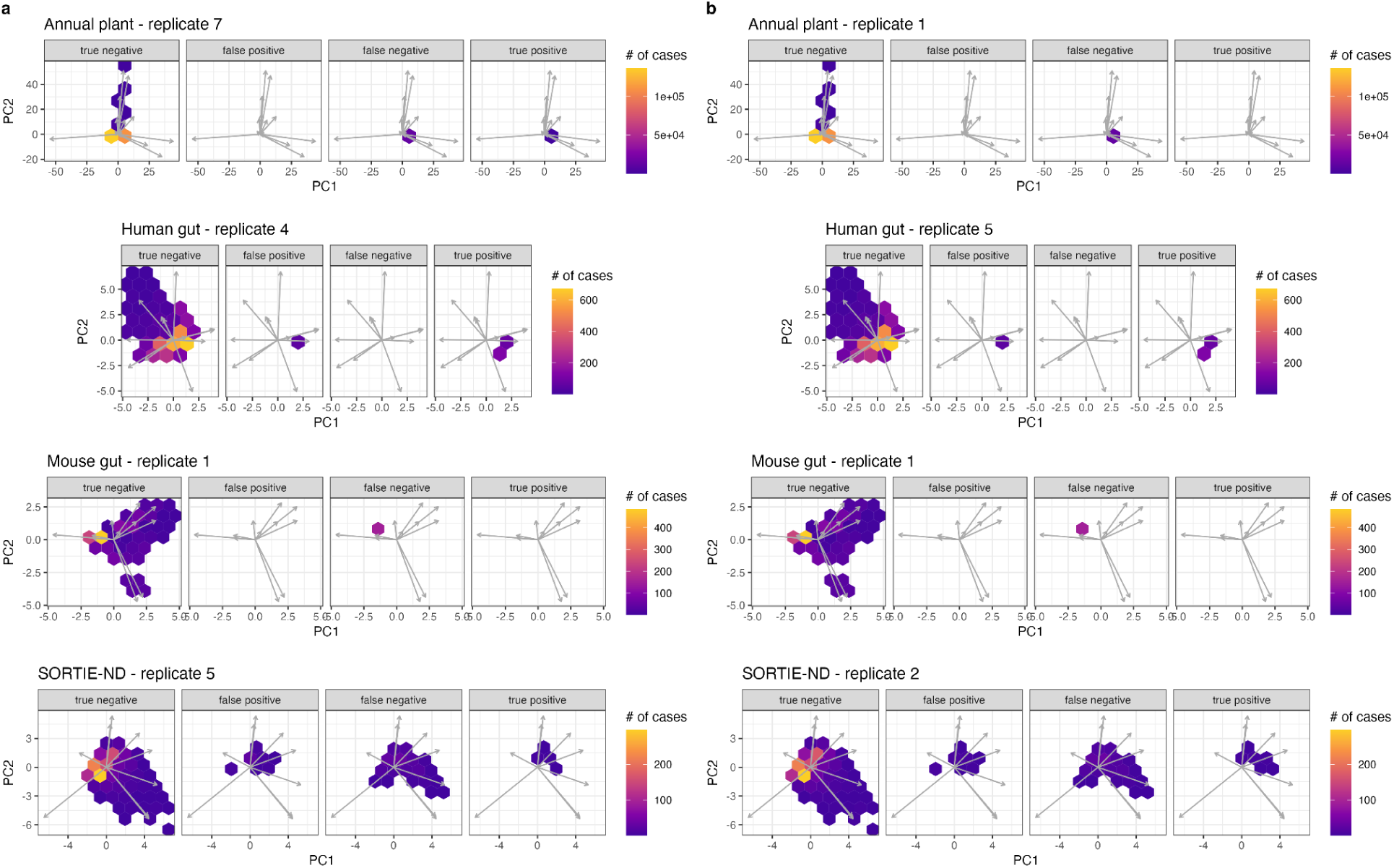
Structure of error types in abundance outcome space for the maximizing total abundance task for each dataset. Prioritizations are for a random forest method, a mixed richness experimental design, and **(a)** 89 or **(b)** 264 training experiments. In each panel, arrows show variable loadings of a principal component analysis in outcome abundance space (*x*^1/4^ transformed to reduce outlier effects); hexagon colors indicate numbers of outcomes that fall within each bin. Outcomes are grouped by whether the experiment yielding them is a true positive, true negative, false positive, or false negative with respect to the prioritization task.

## Footnotes

1 These files will be archived upon acceptance at Dryad or a similar repository.

